# Engineered paediatric tumours retain maintains tumour genotype and phenotype for precision medicine

**DOI:** 10.1101/2024.11.17.619539

**Authors:** MoonSun Jung, Valentina Poltavets, Joanna N. Skhinas, Gabor Tax, Alvin Kamili, Angela (Jinhan) Xie, Sarah Ghamrawi, Philipp Graber, Jie Mao, Marie Wong-Erasmus, Louise Cui, Kathleen Kimpton, Pooja Venkat, Chelsea Mayoh, Emmy D. G. Fleuren, Ashleigh M. Fordham, Zara Barger, John Grady, David M. Thomas, Eric Yiwei Du, Mark J. Cowley, Andrew J. Gifford, Jamie I. Fletcher, Loretta M. S. Lau, M. Emmy M. Dolman, J. Justin Gooding, Maria Kavallaris

**Affiliations:** Children’s Cancer Institute, Lowy Cancer Research Centre, UNSW Sydney, Sydney, New South Wales, 2052, Australia; NHMRC Clinical Trials Centre, Faculty of Medicine and Health, The University of Sydney, Sydney, New South Wales, 2050, Australia; School of Clinical Medicine, UNSW Medicine & Health, UNSW Sydney, Sydney, New South Wales, 2052, Australia; Australian Centre for NanoMedicine, UNSW Sydney, Sydney, New South Wales 2052, Australia; Melanoma Institute Australia, New South Wales, 2065, Australia; The Garvan Institute of Medical Research and The Kinghorn Cancer Centre, Darlinghurst, New South Wales, 2010, Australia; School of Chemistry, UNSW, Sydney, New South Wales, 2052, Australia; Anatomical Pathology, NSW Health Pathology, Prince of Wales Hospital, Randwick, New South Wales, Australia; Kids Cancer Centre, Sydney Children’s Hospital, Sydney, New South Wales, 2031, Australia; Princess Maxima Center for Paediatric Oncology, 3584 CS Utrecht, The Netherlands

**Keywords:** paediatric cancers, preclinical models, tumour organoid models, 3D bioprinting, high-throughput (HTP) drug screening

## Abstract

Precision medicine for paediatric and adult cancers that includes drug sensitivity profiling, can identify effective therapies for individual patients. However, obtaining adequate biopsy samples for high-throughput (HTP) screening remains challenging, with tumours needing to be expanded in culture or patient-derived xenografts – this is time-consuming and often unsuccessful. Herein, we have developed paediatric patient-derived tumour models using an engineered extracellular matrix (ECM) tissue mimic hydrogel system and HTP 3D bioprinting. Gene expression analysis from neuroblastoma and sarcoma patients identified key components of the ECM in these tumour types. Engineered hydrogels with ECM-mimic peptides were used to create patient-specific tumour organoids, modelling tumour growth conditions. Expanded tumour organoids recapitulated the genetic and phenotypic characteristics of the original tumours and retained tumourgenicity. Screening of these models identified individualised drug sensitivities. Our approach offers a timely and clinically relevant technology platform for precision medicine in paediatric cancers, potentially transforming preclinical testing across cancer types.

## Introduction

Cancer remains the leading cause of disease-related death in children in developed countries^1^. High-risk cancer makes up 30% of all cases and prognosis is poor despite patients receiving intensive treatments that cause significant side effects and dose-limiting toxicities^2^. Neuroblastoma is the most common extracranial solid tumour, with overall survival rate of ∼60% for high-risk disease^3^. Sarcomas are heterogeneous mesenchymal tumours arising from bone or soft tissue, with patients having metastatic disease showing overall survival rates of 30% for Ewing sarcoma^4^ and 45% for osteosarcoma^5^, respectively. Neuroblastoma and sarcoma patients with high-risk disease have limited therapeutic options. Moreover, many of the therapies given to treat patients with recurrent and drug-resistant disease are highly toxic, which means that children can be exposed to damaging and ineffective therapies. Indeed, survivors have a high likelihood of experiencing life-long health issues^6^.

With advancement of sequencing technologies, molecular-based precision medicine has gained momentum in cancer management to identify actionable mutations and associated treatments for paediatric patients^7–10^. However, genomic testing alone does not benefit all patients, since ∼30% of paediatric patients do not have actionable molecular alterations^11^ and even then, targeting an actionable alteration with a specific drug results in objective clinical response in the minority of patients^9,11^. Tumour drug sensitivity profiling (DSP) holds promise to support clinical treatment decision-making in precision medicine and expand therapeutic options for cancer patients^8,12,13^. In precision medicine platforms, tumour biopsies often lack sufficient material for use beyond routine diagnostic histopathology and genomic analysis, precluding comprehensive DSP. Subsequent cell expansion is often required *via* primary cell culture or the development of patient-derived xenograft models (PDXs) ^14–16^. Among the most common challenges with expansion of primary material are that not all samples are able to grow in culture or successfully engraft *in vivo.* Furthermore, time required for in vitro expansion of patient material using PDXs is highly variable (up to 12 months) ^8,13,17,18^.

*Ex vivo* tumour organoid models – cancer cells grown in three-dimensional (3D) structures – are increasingly being used to test drug responses of both adult^19,20^ and paediatric^21,22^ patient tumours. Tumour organoid models that are representative of the patient tumour can be expanded in a robust manner, are reflective of tumour heterogeneity, and amenable to rapid DSP and mechanistic studies will be transformative for precision medicine. However, the ability to grow tumour cells as organoids directly from biopsies is highly variable and cancer type dependent^16,23^. For example, osteosarcomas are difficult to grow *ex vivo*, often requiring *in vivo* engraftment to expand cell numbers for drug screening ^24,25^. Furthermore, there is a lack of high-throughput (HTP) platforms that efficiently support *ex vivo* growth of patient tumour cells and are also compatible with HTP drug screening^26,27^. Moreover, many *ex vivo* models of cancers lack consideration of the extracellular matrix (ECM) which can impact cellular signalling and response to therapy ^28,29^. Most 3D models that attempt to incorporate the ECM are grown in an animal-derived matrix which suffers batch-to-batch variability that impacts cellular cues and stiffness^30,31^. Mimicking relevant physiological components of the tissue extracellular environment, including ECM protein components and matrix stiffness, will be an important advance in precision medicine.

We have recently developed a 3D bioprinting platform whereby cultured cells are embedded in synthetic hydrogels with tunable properties that can mimic a range of ECM components and spatial growth constraints of a tumour^32,33^. This platform allows rapid and reproducible encapsulation of cells with high viability and enables the expansion of a range of cell types in ECM-mimic hydrogels^34^. A key question was whether this approach could be developed to support the growth and expansion of patient tumour cells for precision medicine. Here, we investigated the potential of 3D bioprinting and this ECM-mimic hydrogel system to establish high-risk neuroblastoma and sarcoma patient-derived tumour organoids that maintain the genomic and phenotypic characteristics of the patient tumours and incorporate these organoids into HTP drug testing for precision cancer medicine.

In this proof-of-concept study, we address some key limitations in precision medicine, namely the ability to maintain and expand difficult to grow freshly isolated patient-derived cells in a HTP fashion and conduct robust HTP drug screening in an environment that mimics tumour growth in a timely manner. Our approach holds enormous potential to identify drug sensitivities in cancers and expand therapeutic options for high-risk cancer patients.

## Results

### Defining the extracellular matrix environment of high-risk neuroblastoma and sarcoma tumours

To investigate the extracellular environment conditions for 3D culture of high-risk neuroblastoma and sarcoma patient-derived cells, we analysed tumour-specific expression patterns of ECM genes using recently published methodology^35^ and ECM Organization gene set^36^. RNA sequencing (RNA-seq) data for high-risk paediatric tumours available through the Australian precision medicine program, ZERO^7^, was used to perform bioinformatic analysis of the ECM Organization gene set (n=265) in a cohort of neuroblastoma (NBL) and sarcoma, comprising Ewing sarcoma (EWS), osteosarcoma (OST) and rhabdomyosarcoma (RMS) (n=146) **(Figure 1A)**. The top 30 of 265 structural ECM genes and their regulators highly expressed across the four cancer types were identified **(Figure 1B)**. Many of these are translated to collagens^37^, the most abundant proteins in the human body and fibronectin^38^, abundant in highly organized structures in both interstitial matrix and basement membrane. Further analysis of the core matrisome genes (encoding structural components of the ECM) ^39,40^ confirmed significant upregulation of *FN1* **(Figure 1C)** and *COL1A1* **(Figure 1D)** relative to other genes in the cohort. Thus, our analysis identified key genes that encode for structural components within childhood neuroblastoma and sarcoma tumours. This knowledge was incorporated in determining the 3D extracellular mimic environment to support the growth of neuroblastoma and sarcoma patient-derived cells.

**Figure 1.**
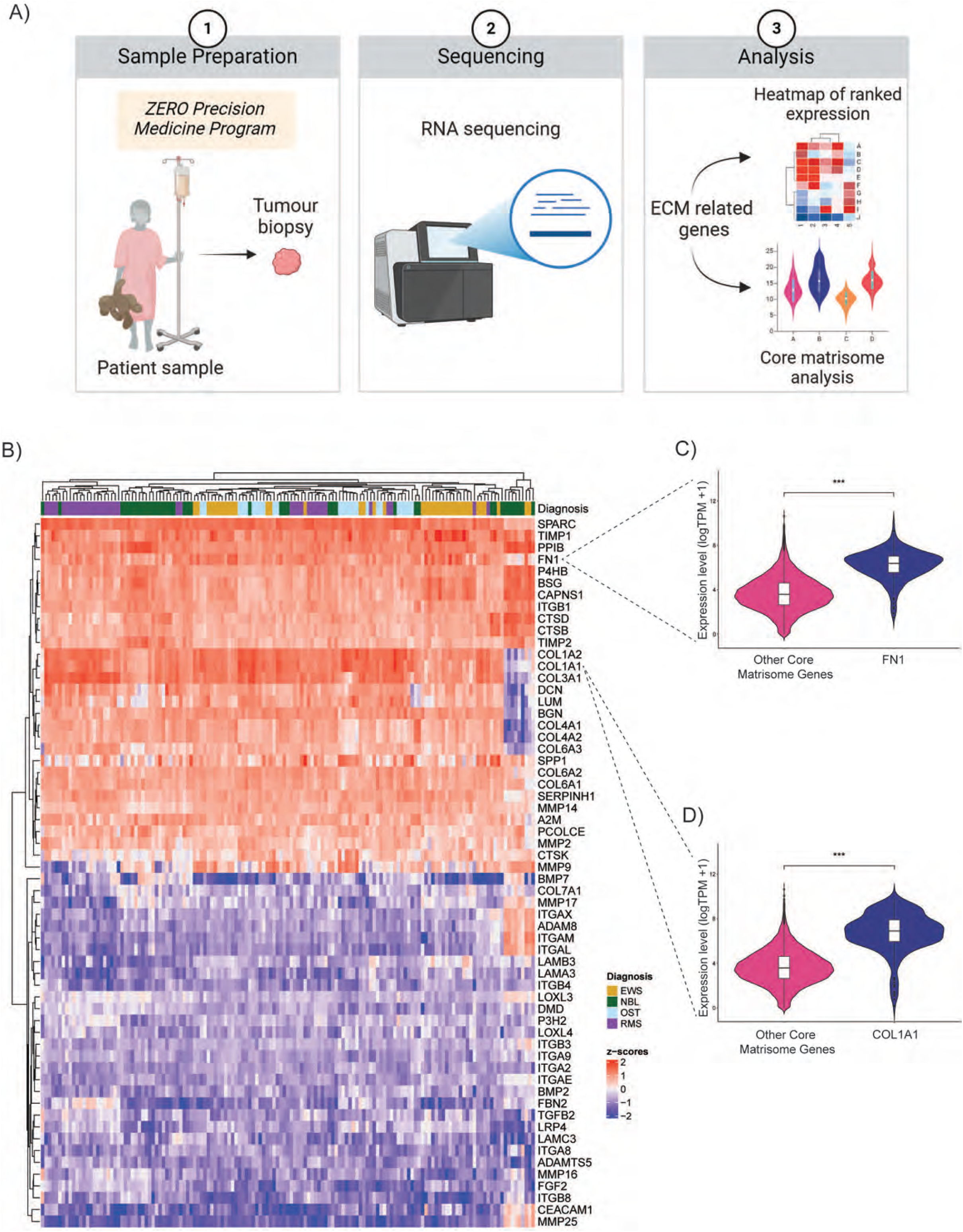
Defining extracellular matrix environment of high-risk neuroblastoma and sarcoma tumours. A) A workflow schematic of generating RNA sequencing results from the paediatric patient samples and subsequent ECM-related gene analysis. B) Heatmap of the top and bottom 30 genes ranked by expression levels out of 265 ECM-relevant genes identified in neuroblastoma and sarcoma samples. Gene TPM values are represented by colour from red (high Z-score, high expression) to blue (low Z-score, low expression). Diagnosis EWS (Ewing sarcoma N=40); NBL (neuroblastoma N=41); OST (osteosarcoma N=25); RMS (Rhabdomyosarcoma n=40). C)-D) Violin plots of collagen type I alpha 1 (*COL1A1*) and fibronectin (*FN1*) expression and compared to the remaining core matrisome ECM genes (n=79) in the cohort. Expression levels are log transcript per million (TPM)+1. Welch’s t-test, P< 0.001

### Tunable ECM-mimic hydrogels support paediatric tumour growth

To grow and expand patient-derived samples, current methods that rely on cell culture or establishment of patient-derived xenografts have limited success^13^. We have obtained eight samples representing neuroblastomas and sarcomas **(Table S1)** that exhibited variable success in generating cell cultures **(Table S2)** and patient-derived xenografts **(Table S3)**.

To systematically assess the mechanical and biological requirements for optimal cellular growth across our tumour types, we tested the capacity of distinct ECM-mimic hydrogel combinations to support tumour growth *in situ*. To create a tumour-like ECM for high-risk neuroblastoma and sarcoma, we selected hydrogels with two different stiffnesses levels - 1.1 kPa and 3 kPa - to closely mimic lung (∼0.8 kPa) ^41,42^ and liver (∼3 kPa) ^43,44^ tissues, respectively. These organs are metastatic sites for neuroblastoma^45^ and sarcomas^46^. We prioritized the selection of fibronectin and collagen I based on the top matrisome genes identified in neuroblastoma and sarcoma patient samples **(Figure 1B).** Furthermore, we expanded our functional peptide selection with the inclusion of laminin in the hydrogels. Laminin is indispensable for integrin-mediated cell adhesion and a major component of both the basement membrane and Matrigel^47^. The hydrogels were engineered to incorporate an integrin-binding peptide of fibronectin (RGD)^48^, a collagen I sequence (GFOGER)^49^, and a laminin peptide fragment (DYIGSR)^50^. We tested a total of six hydrogel conditions incorporating either fibronectin (FN) alone, a combination of fibronectin and collagen I (FN + CN), or a tripeptide combination of fibronectin, collagen I and laminin (FN + CN + LN), at a stiffness of either 1.1 kPa or 3 kPa.

Patient-derived cells from eight high-risk paediatric tumours were bioprinted in the six hydrogel conditions and their proliferative capacity measured by AlamarBlue assay over a 14-day period. Post-bioprinting, patient-derived neuroblastoma **(Figure 2Ai)**, Ewing sarcoma **(Figure 2Bi)** and osteosarcoma **(Figure 2Ci)** cells embedded in the tripeptide hydrogel (FN + CN + LN) exhibited similar growth dynamics, characterized by consistent proliferative activity up to day 14. Cells remained highly viable for 14 days of culture in both the 1.1 kPa **(Figure 2A-Cii)** and 3 kPa **(S2A-C)** tripeptide hydrogel. Additionally, we observed similar growth dynamics in one peptide (FN) and two peptide (FN + CN) hydrogels at 1.1kPa and 3kPa stiffness **(S1A-C).**

**Figure 2.**
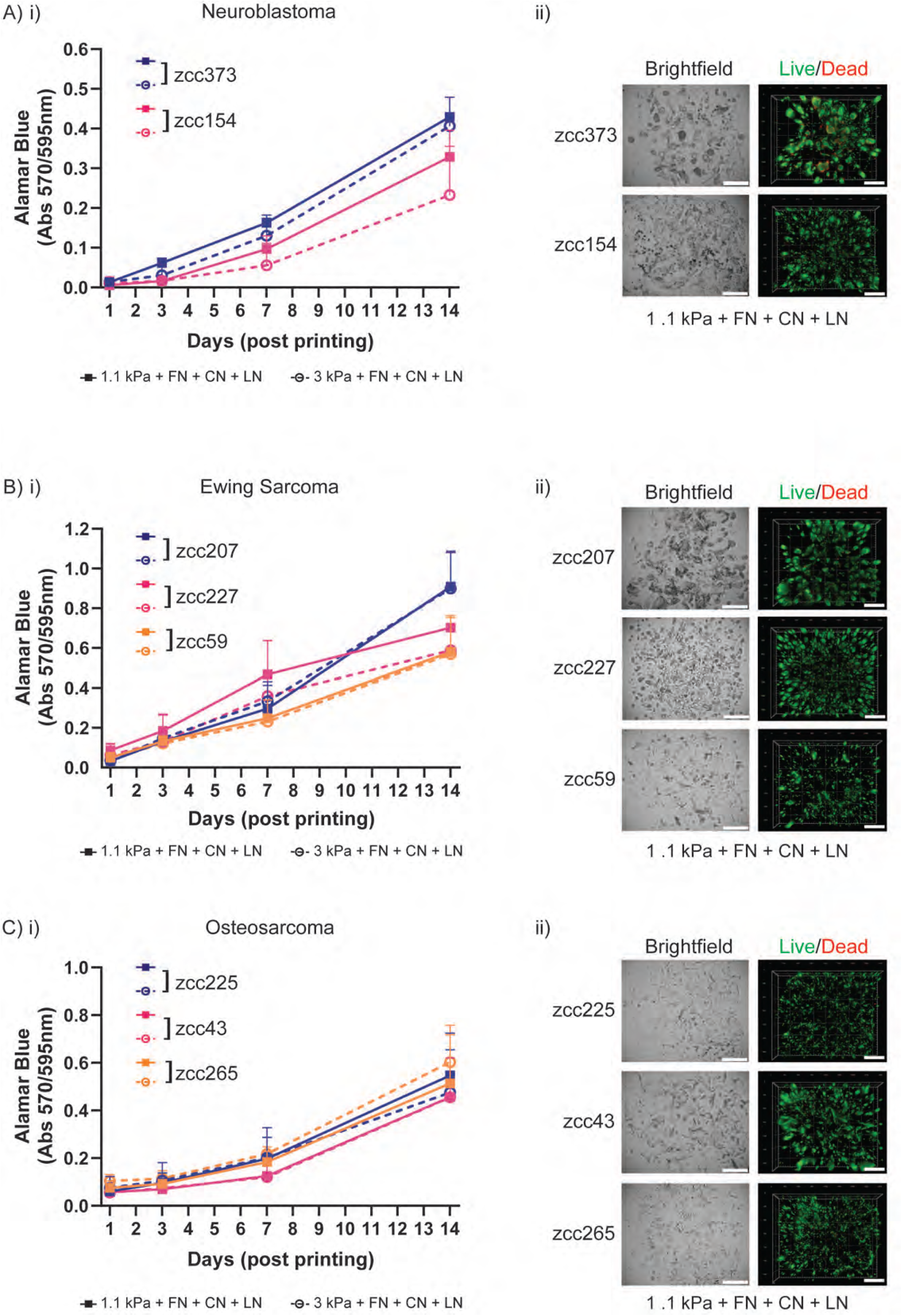
Tunable ECM-mimic hydrogels support paediatric tumour growth. A)-C) i) Cell proliferation of 3D bioprinted patient-derived cells grouped by disease type: neuroblastoma (zcc373, zcc154), Ewing sarcoma (zcc207, zcc227, zcc59) and osteosarcoma (zcc225, zcc43, zcc265). Each sample was bioprinted in either 1.1kPa or 3kPa PEG hydrogels, both containing FN (fibronectin), CN (collagen 1) and LN (laminin) peptides. Proliferation rate was measured at day 1, 3, 7 and 14 post-printing. Data are representative of mean ±SD. All experiments were repeated three times. A)-C) ii) Viability of 3D bioprinted patient samples in 1.1kPa gels containing FN, CN and LN peptides, grouped by cancer type. Cells were stained with calcein-AM (green; live)/ethidium homodimer 1 (red; dead) Live/Dead Assay. Z stack 3D images were taken at day 14 post-printing. Representative images shown for each patient sample in brightfield (left) and Live/Dead (right). Scale bars on all images are 500 µm.

Our analysis examined the influence of hydrogel stiffness and incorporation of functional peptides on cellular growth. Collectively our data suggest that neither the stiffness, nor specific peptide combinations had a major impact on cellular proliferation across all disease types **(Figure 2, S1 and S2).** However, given the pivotal roles of cell adhesion peptides and the basement membrane for cellular function, along with our findings in the paediatric tumour ECM associated genes **(Figure 1C-D)**, we strategically selected the tripeptide combination of fibronectin, collagen I and laminin-mimicking peptides at 1.1 kPa stiffness for subsequent experiments. This hydrogel was selected for subsequent experiments as it supports the cellular growth of high-risk paediatric tumour cells across multiple cancer types

### 3D bioprinted tumour organoids retain primary molecular, phenotypic and tumourigenic characteristics

Having identified the ECM-mimic hydrogel conditions supportive of high-risk paediatric neuroblastoma and sarcoma cell growth *ex vivo*, we systematically investigated the similarity between the 3D bioprinted tumour models and the original tumours. Patient-derived cells from the eight high-risk paediatric tumours were bioprinted, cultured for seven to 14 days and retrieved from the hydrogels for targeted capture sequencing of 523 pan-cancer genes (Illumina TruSight Oncology 500) **(Figure 3A)**. We compared key molecular characteristics between 3D bioprinted tumour organoids and original patient samples previously identified by whole genome sequencing (WGS) and RNA-seq by ZERO ^7^.

**Figure 3.**
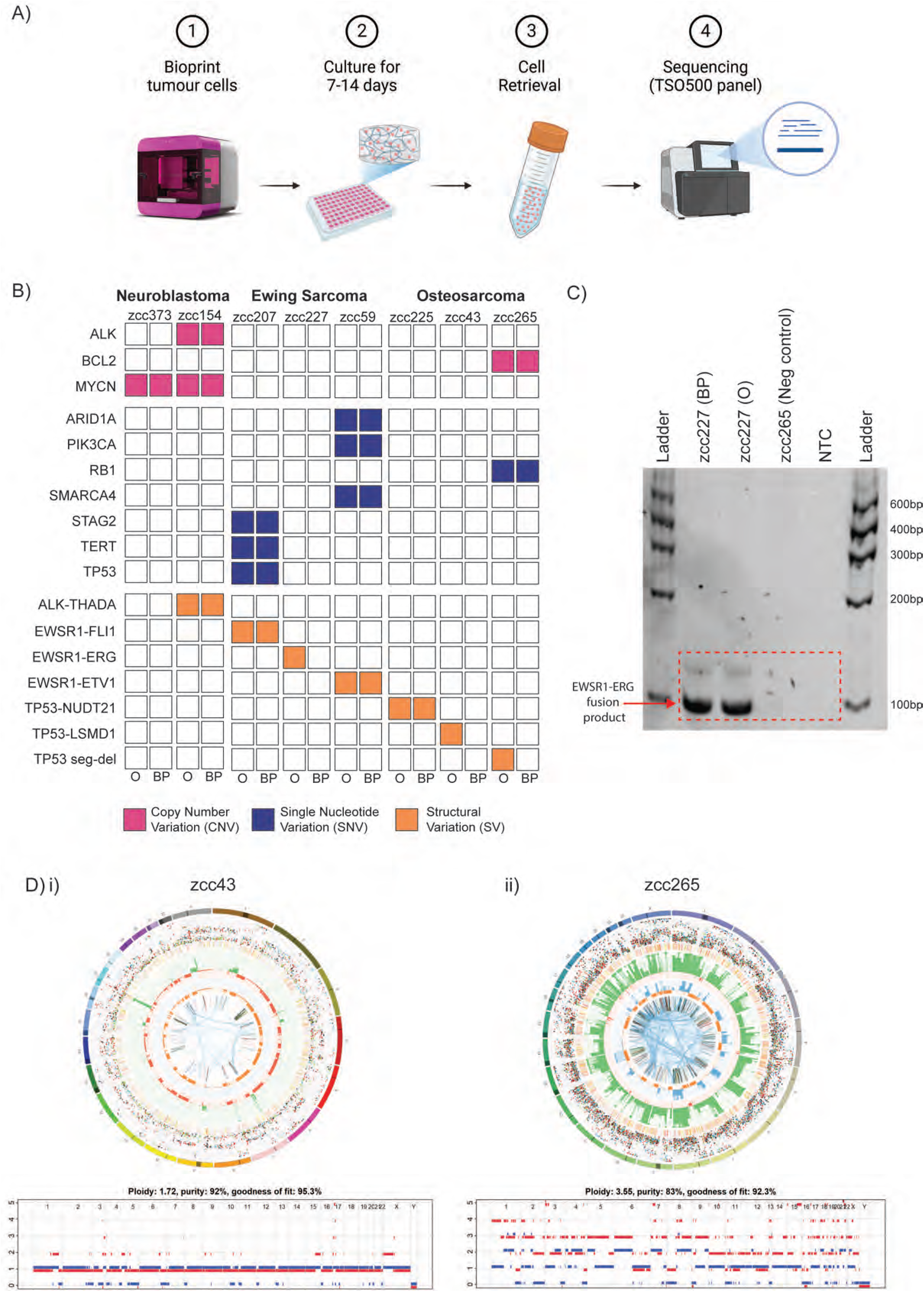
3D bioprinted tumour organoids retain primary molecular characteristics. A) Schematic showing workflow of obtaining bioprinted patient-derived samples for sequencing. Patient-derived samples were bioprinted in 1.1kPa + FN + CN + LN hydrogels and cultured for 7 to14 days, then cells were retrieved and purified for sequencing against the TruSight Oncology 500 panel. B) Reportable pathogenic events for each patient-derived sample in either the Original (O) sample or the Bioprinted (BP) sample, colour coded by mutation type. C) PCR analysis of the DNA from bioprinted sample zcc227 (BP) and original patient zcc227 (O) confirming the presence of *EWSR1-ERG* structural variant. Zcc265 (osteosarcoma sample) used as negative control; NTC – no template control; D) i-ii) Circos plots showing genome-wide profile of the original zcc43 and zcc265 osteosarcoma patient tumours identified through whole genome sequencing (WGS) in comparison with ASCAT copy number profiles for bioprinted samples generated from Single Nucleotide Profiling (SNP).

Overall, we observed that the molecular drivers of bioprinted patient-derived cells recapitulated those found in the original patient samples, although the clonal composition of some samples varied slightly. High-risk neuroblastoma bioprinted tumour organoids retained disease-specific driver alterations such as *MYCN* (zcc373, zcc154) and *ALK* amplification (zcc154) (**Figure 3B**, **Table S4**). Ewing sarcoma tumour organoids preserved characteristic *EWSR1* rearrangements with erythroblast transformation-specific (ETS) family of transcription factors such as *FLI1*, *ERG* and *ETV1*^51^. Moreover, Ewing sarcoma sample zcc207 retained *STAG2* (p.Leu1132Ter) nonsense variant, *TERT* (c.-57A>C, 5’ UTR) noncoding single nucleotide variant and *TP53* (p.His193Tyr) missense variant (**Figure 3B, Table S4**). Sample zcc59 preserved *ARID1A* (p.Arg1287LysfsTer11) nonsense variant and *SMARCA4* (p.Arg1157Trp) missense variant. Overall, the tumour-specific subclones were preserved in the bioprinted tumour organoids. We observed some clonal enrichment where variant allele frequency (VAF) was higher in zcc59 *(SMARCA4)* and zcc207 (*PIK3CA)* compared to original patient samples **(Table S4).** In the case of Ewing sarcoma zcc227 bioprinted sample, targeted RNA sequencing did not identify the *EWSR1-ERG* fusion. However, subsequent PCR analysis confirmed the presence of 100 bp amplicon corresponding to the *EWSR1-ERG* fusion breakpoint in both the original patient DNA and the bioprinted sample (**Figure 3C**).

All three osteosarcoma tumours - zcc225, zcc43, and zcc265 - harboured structural variations in *TP53*. Targeted sequencing confirmed the preservation of the *TP53-NUDT1* fusion in the matching bioprinted organoid for zcc225 **(Figure 3B).** Original zcc43 sample contained intragenic structural variant of *TP53* with breakpoint in the noncoding region (exon1), and zcc265 sample included intergenic *TP53-LSMD1* structural variant. These types of SVs are known to be very challenging to detect using targeted DNA sequencing assays^52^, we subsequently confirmed the genetic match with amplicon analysis. Bioprinted tumour organoids zcc43 and zcc265 were further characterized using both short tandem repeat (STR) and single-nucleotide polymorphism (SNP) profiling. STR profiles of bioprinted samples were 100% identity match to the original patient tumour. Additionally, SNP profiling confirmed high percentages of tumour cell content in the bioprinted tumour organoids with overall ploidies and copy number profiles matching the original patient samples (**Figure 3Di** and **3Dii**). For zcc43, the tumour cell content was 92% with an overall ploidy of 1.72, compared to 86% and an overall ploidy of 1.64 in the original sample (**Figure 3Di**). Similarly, zcc265 exhibited a tumour cell content of 83% with an overall ploidy of 3.55, compared to 61% and an overall ploidy of 3.93 in the original sample (**Figure 3Dii**). Moreover, in osteosarcoma bioprinted sample zcc265, targeted sequencing confirmed the presence of *RB1* (p.Glu170ValfsTer6) nonsense variant and *BCL2* amplification **(Figure 3B)**. Taken together, our data indicates that 3D bioprinted tumour organoids retained critical pathogenic DNA variants and key molecular characteristics, confirming their close resemblance to the original tumours.

We next characterized the morphologic features of the 3D bioprinted tumour organoids in comparison with the features of the patient-derived xenografts (PDXs) from which they were derived. Patient-derived cells were bioprinted and cultured for 7 to 14 days, prior to preparation and cryosectioning. Samples were stained with both routine H&E and immunohistochemical staining. The bioprinted tumour organoids recapitulated the morphologic features of the PDX from which they were derived **(Figure 4A-Ci)**. Each of the bioprinted tumour organoids contained “small round blue cells” in H&E staining, consisting of tumour cells with round to oval hyperchromatic nuclei and minimal cytoplasm **(Figure 4A-Cii)**. The proliferative capacity of the bioprinted cells was demonstrated by a high proportion of cells expressing Ki-67 **(Figure 4A-Ciii)**. The tumour type was confirmed by diagnostic immunohistochemical staining. The neuroblastoma tumour organoids demonstrated diffuse strong nuclear PHOX2B^53,54^ staining **(Figure 4Aiv)**, Ewing sarcoma tumour organoids circumferential membrane staining for CD99^55,56^ **(Figure 4Biv)**, and osteosarcoma tumour organoids with nuclear SATB2^57,58^ staining **(Figure 4Civ)**. These results confirm that the morphologic and diagnostic immunohistochemical characteristics of the original patient samples are preserved in patient-derived 3D bioprinted tumour organoids. Lastly, to ensure phenotypic features of the original cells were maintained, we examined tumourigenic capacity of patient-derived cells after 3D bioprinting and expansion. Two tumour types were examined for tumourigenic potential post bioprinting: one neuroblastoma (zcc373) and one Ewing sarcoma (zcc207). Patient-derived cells dissociated from the bioprinted tumour organoids were subcutaneously engrafted into immunocompromised NSG mice (n=4 per group), while non-bioprinted *in vivo* expanded cells were used as a control **(Figure 5A)**. Both neuroblastoma (zcc373) and Ewing sarcoma (zcc207) tumour cell growth *in vivo* was comparable before and after bioprinting (**Figure 5Bi and Ci**). Moreover, tumour morphology, Ki-67 proliferative index and tumour immunohistochemical staining for PHOX2B (neuroblastoma) and CD99 (Ewing sarcoma) remained unchanged between tumours generated from non-bioprinted and bioprinted cells (**Figure 5Bii and Cii**). Taken together, these analyses confirm that 3D bioprinting conditions for high-throughput generation of tumour organoids from neuroblastoma and sarcoma patient-derived cells do not alter the *in vivo* tumourigenic capacity of the cells.

**Figure 4.**
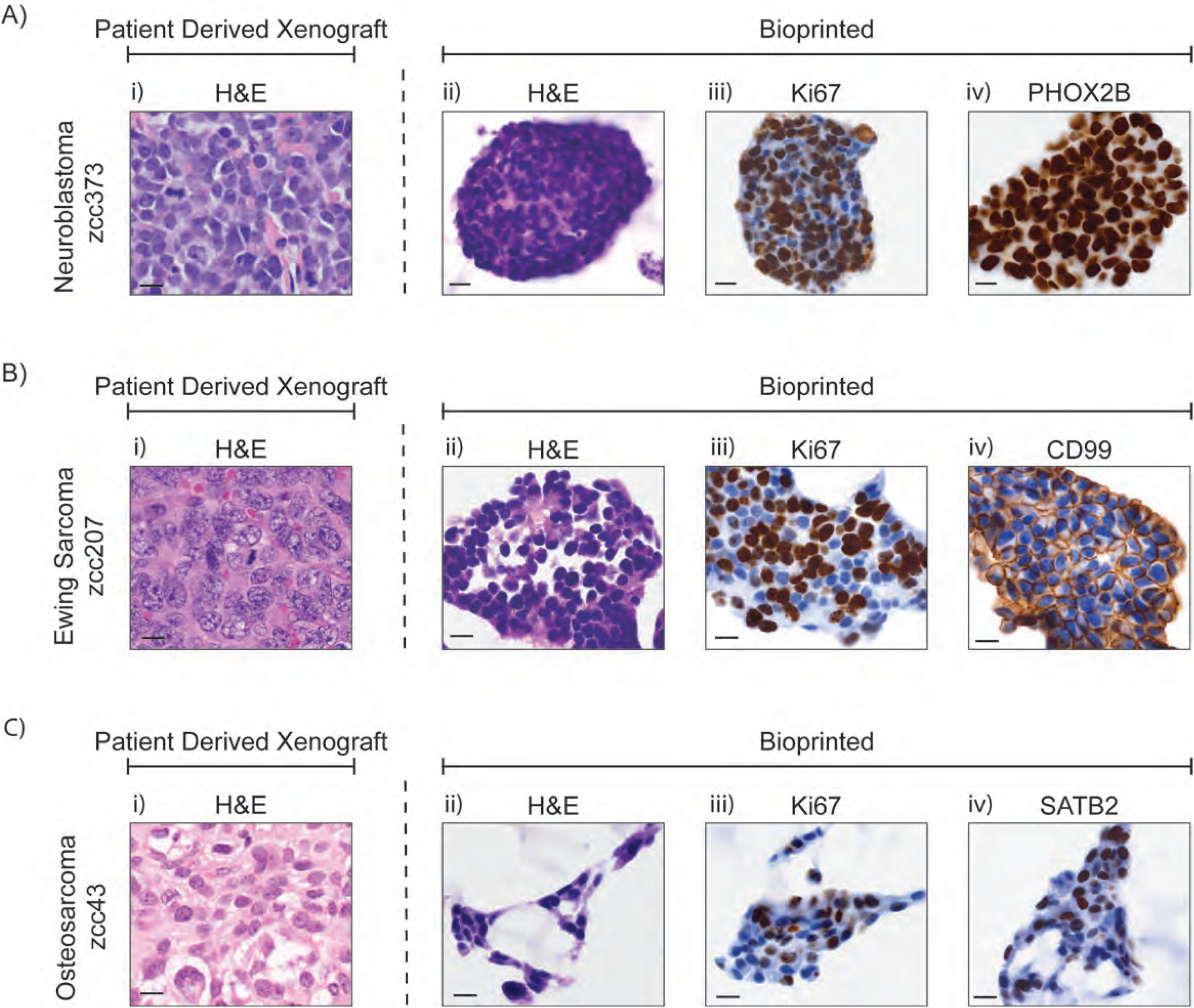
3D bioprinted tumour organoids retain disease-specific phenotypic characteristics and proliferative capacity in situ. A)-C) i) Representative histology images of original patient-derived xenografts from neuroblastoma (zcc373), Ewing sarcoma (zcc207) and osteosarcoma (zcc43) paediatric patients (left panels). A)-C) ii) Representative histology and iii-iv) immunohistochemistry images of bioprinted patient-derived cells stained with H&E (left), proliferation marker Ki-67 (middle) and tumour-specific markers (right) PHOX2B for neuroblastoma, CD99 for Ewing sarcoma and SAT2B for osteosarcoma. Scale bars are 10 µm for all images.

**Figure 5.**
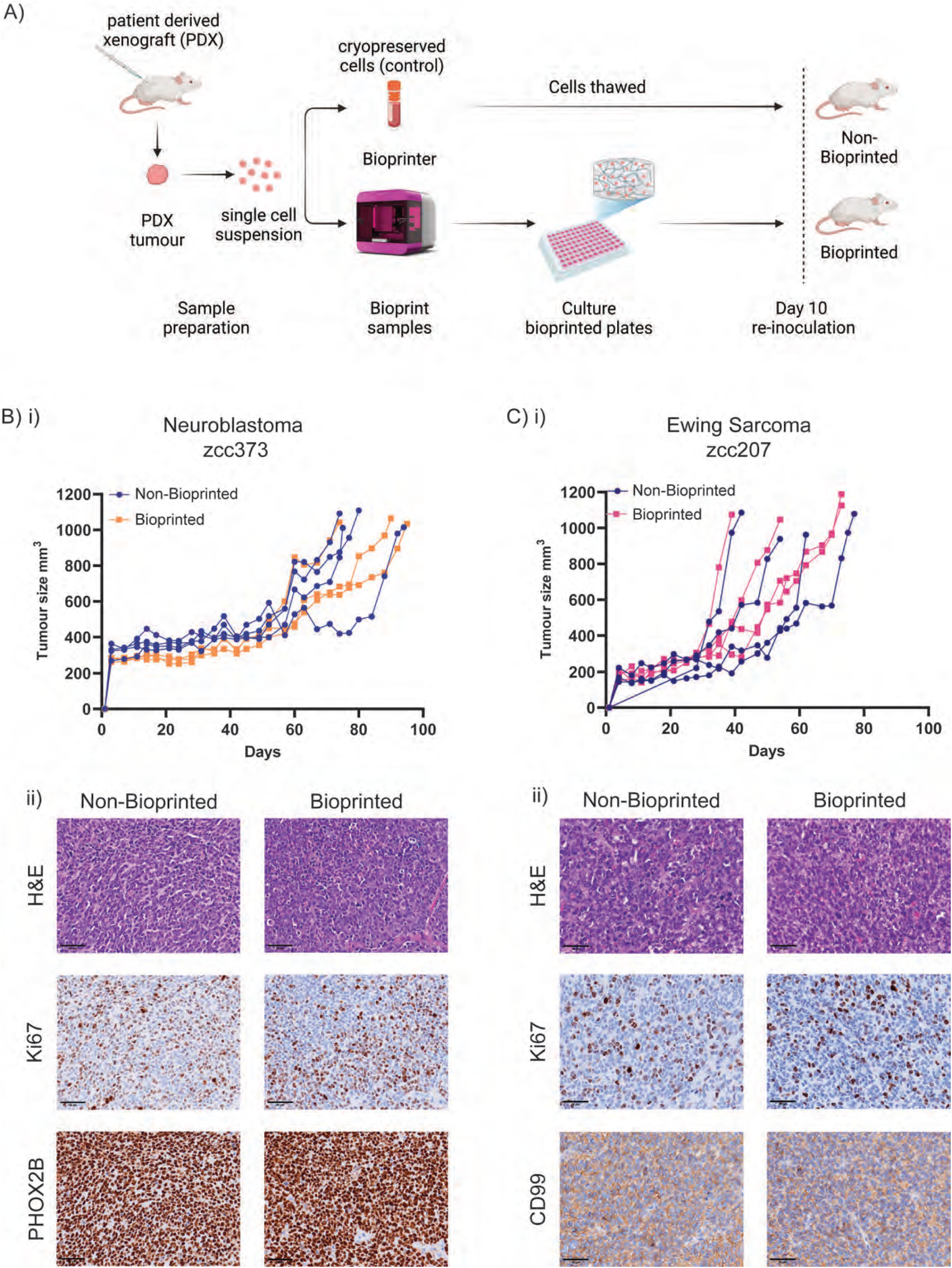
Bioprinted patient-derived cells retain their tumourigenicity *in vivo*. A) Schematic workflow of in vivo tumourigenicity experiment. Briefly, one neuroblastoma and one Ewing sarcoma patient-derived xenograft tumour were dissociated to single cell suspension and bioprinted in 1.1kPa + FN + CN + LN hydrogels for 10 days. At day 10, cells were retrieved and re-inoculated into NSG mice, with cryopreserved cells from the same tumour acting as control non-bioprinted samples. N=4 mice per condition. B) and C) i) Individual tumour growth curves for neuroblastoma and Ewing sarcoma tumours respectively. One mouse was excluded from zcc373 Bioprinted condition due to external injury. B) and C) ii) Representative histology and immunohistochemistry images from tumours derived from non-bioprinted (left) and bioprinted (right) samples stained with H&E (top row), proliferative marker Ki-67 (middle row) and tumour-specific markers PHOX2B for neuroblastoma or CD99 for Ewing sarcoma (bottom row). Scale bars for all images are 50 µm.

### 3D bioprinted tumour organoids are compatible with a high-throughput preclinical drug screening platform and reveal patient-specific drug vulnerabilities

Having confirmed our ability to grow and expand patient-derived cells in an ECM-mimic environment while preserving their key molecular and phenotypic characteristics *ex vivo*, we proceeded to assess the compatibility and feasibility of 3D bioprinted tumour organoids with the HTP *ex vivo* drug screening platform established by ZERO ^13^. We designed a custom-made drug library of 48 compounds selected from US Federal Drug Administration (FDA)-approved standard of care drugs for neuroblastoma and sarcoma patients, therapeutic agents previously identified by ZERO to be effective in specific high-risk paediatric samples^15^ and some previously unconsidered targets of potential interest **(Table S5).** Patient-derived cells were bioprinted, cultured between 7 to 14 days and subjected to an established ZERO pipeline for *ex vivo* drug screening using our customized library **(Figure 6A)**.

**Figure 6.**
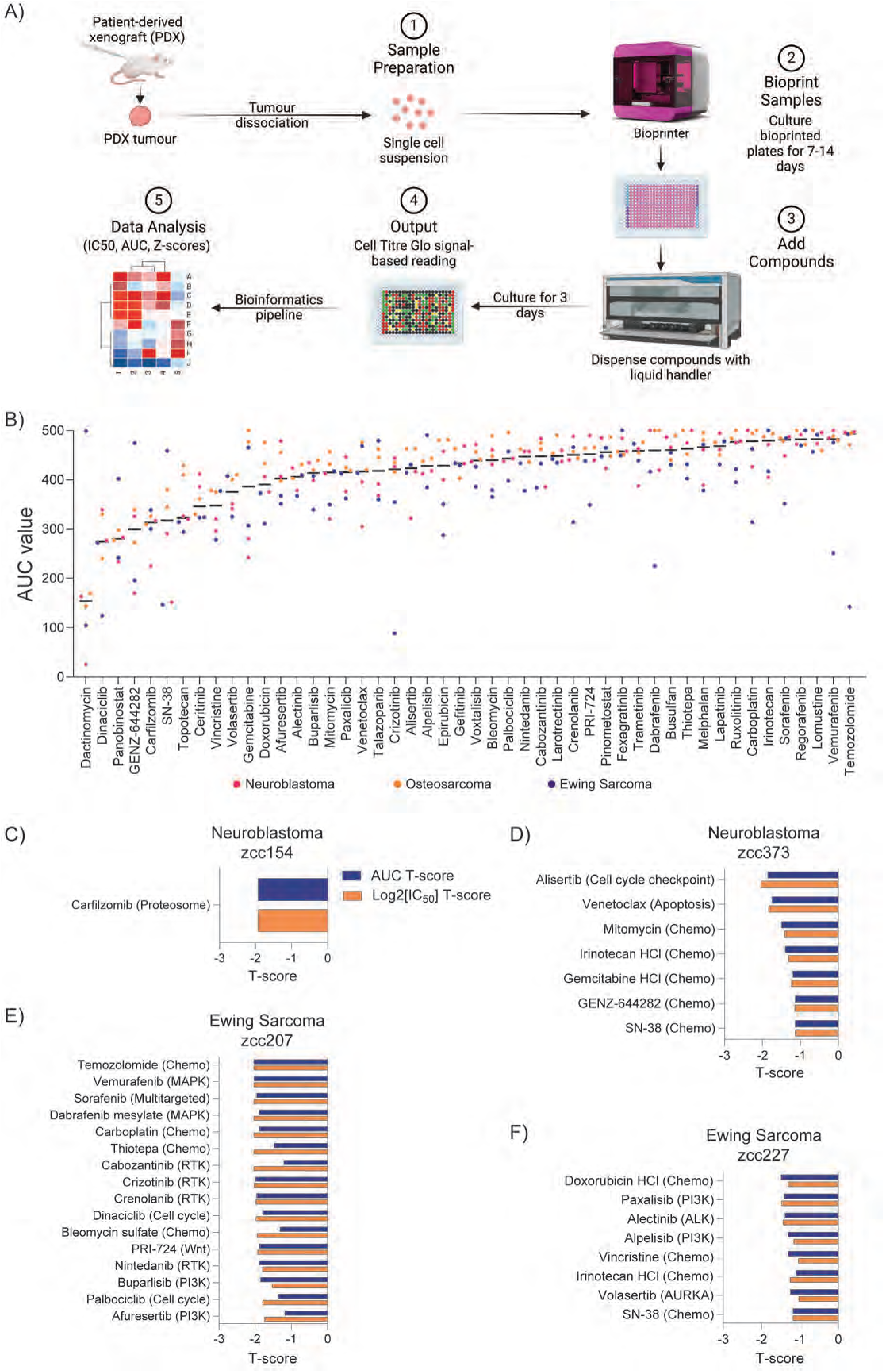
3D bioprinted tumour organoids are compatible with a high-throughput preclinical drug screening platform and reveal patient-specific drug vulnerabilities. A) Schematic showing HTP drug screening pipeline. B) AUC (area under the curve) distribution across all samples for custom drug library of 48 compounds. Drugs are ordered from most (left) to least (right) effective based on lowest median AUC. Colours indicate tumour type (pink – neuroblastoma, orange – osteosarcoma, blue – Ewing sarcoma) C)-F) Bar graphs representing tumour-specific drug sensitivity to drugs based on the combined AUC (blue) and log2 [IC50] (orange) T-scores for each drug. T-score ≤1 for both values considered a differential response.

High-throughput drug screening was performed against six patient-derived samples. To examine overall agent class susceptibility in our samples, we ranked the drugs by lowest median area under the curve (AUC) and log2 [IC50] values. From the analysis, we found that the most potent compounds for the six high-risk cancer patients were dactinomycin (RNA synthesis inhibitor); panobinostat (pan-HDAC inhibitor); dinaciclib (CDK1/2/5/9 inhibitor); SN-38 and GENZ-644282 (DNA topoisomerase I inhibitors); carfilzomib (proteasome inhibitor) and ceritinib (ALK inhibitor) (**Figure 6B**, Figure **S3**). This is consistent with findings across multiple paediatric preclinical drug screen studies which identified general responses across all tumour types to chemotherapeutic agent dactinomycin (actinomycin D) ^8,12^ and targeted agents panobinostat and dinaciclib ^13^, carfilzomib ^12^ and ceritinib ^8^.

To investigate tumour-specific drug sensitivity, we compared T-score values of AUC and log2 [IC50] based on the most differential drug responses in the cohort. We identified tumour-specific drug sensitivity to one or more library compounds (T-score ≤1) in four out of six screened samples. We detected unique differential sensitivities to targeted agents in two neuroblastoma and two Ewing sarcoma samples. Our analysis revealed that neuroblastoma zcc373 was most differentially sensitive to aurora-A kinase (AURKA) inhibitor alisertib and B-cell leukemia/lymphoma 2 (BCL-2) inhibitor venetoclax **(Figure 6D, Table S6)**. Interestingly, this sample did not contain molecular indicators for BCL-2 sensitivity. However, this sample had MYCN amplification **(Figure 3B)**, which could explain the observed sensitivity to both alisertib and venetoclax. Alisertib alters the conformation of AURKA, thereby disrupting its association with MYCN and inducing its ubiquitin-dependent degradation^59^. MYCN amplification primes neuroblastoma cells by upregulating pro-apoptotic BCL-2 family member NOXA, creating vulnerability for BCL-2 inhibition by venetoclax^60^. Interestingly, another MYCN amplified neuroblastoma sample zcc154 **(Figure 3B)** was less sensitive to both alisertib and venetoclax despite containing MYCN amplification **(Table S6)**. This sample was differentially sensitive to proteasome inhibitor carfilzomib (**Figure 6C**). Even though molecular profiling identified ALK amplification **(Figure 3B)** in this sample, it was not differentially sensitive to any of the ALK inhibitors included in our panel (alectinib, crizotinib, ceritinib) **(Table S6),** confirming an established observation in personalized drug testing that the presence of molecular aberrations does not always translate into targeted drug sensitivities *in vitro* ^15^. The zcc207 Ewing sarcoma sample was differentially sensitive to several agents including chemotherapeutics temozolomide, carboplatin and thiotepa. Targeted agents included several receptor tyrosine kinase, MAPK-ERK, PI3K pathway and cell cycle inhibitors **(Figure 6E)**. Ewing sarcoma sample zcc227 showed differential sensitivity to a number of chemotherapeutic agents including doxorubicin, irinotecan and vincristine as well as targeted agents affecting PI3K pathway, receptor tyrosine kinase (ALK) and cell cycle (AURKA) signalling pathways (**Figure 6F).** Osteosarcoma samples zcc225 and zcc265 were generally sensitive to dactinomycin, dinaciclib (CDK1/2/5/9 inhibitor), carfilzomib (proteasome inhibitor) and ceritinib (ALK inhibitor) (**Figure 6B, Figure S3**). Compared to other samples in the cohort, no patient-specific drug sensitivities in osteosarcoma samples were detected. This could be explained in part because many genetic drivers in paediatric osteosarcoma are transcription factors, epigenetic regulators, and fusion oncoproteins, which present significant challenges for targeting ^61^. We suggest that application of a larger and more targeted compound library for osteosarcomas will be beneficial for expanding differential sensitivity readouts for this cancer type. Together our analysis demonstrates the feasibility and compatibility of 3D bioprinted matrix-embedded tumour organoids with a pre-clinical HTP drug screening pipeline as well as the ability to identify differential patient-specific responses and resistance using our workflow.

## Discussion

Precision medicine for childhood cancer that incorporates drug sensitivity profiling (DSP) can identify tailored therapies, avoiding the use of ineffective drugs and toxicity. Herein, we reveal the successful development of high-throughput 3D bioprinted tumour organoid models for high-risk paediatric tumours and their application in personalized DSP. Our models are uniquely engineered to be (i) reflective of the tumour-specific extracellular matrix environment; (ii) representative of the original patient tumours; (iii) expanded in a rapid manner; and (iv) compatible with high-throughput drug screening in a clinically relevant timeframe.

In this proof-of-concept study, we established a 3D bioprinting workflow that enabled robust encapsulation of patient-derived tumour cells in ECM-mimic hydrogels with defined mechanical and biochemical properties. We showed that addressing the tumour extracellular matrix requirements could inform culture conditions required for growth and expansion of non-epithelial tumour cells, which has previously proven to be challenging ^22,25,62^. This is the first study to report incorporating cell-adhesion peptides in the hydrogel systems for paediatric patient-derived cells in a high-throughput manner and apply this workflow in a pre-clinical drug testing pipeline. The hydrogels used in our study are highly tunable, well-defined and display reproducible stiffness and presentation of adhesive cues from batch-to-batch and are designed for use in a HTP 3D bioprinting platform ^63^. This is a significant advantage over the use of animal-derived ECM-gels such as Matrigel which is frequently used in HTP drug screening approaches^64,65^. Matrigel has limitations in accurately replicating tissue-specific stiffness^66^ and has high batch-to-batch variability, thus lacking standardization for HTP drug screening platforms^67^.

In paediatric pre-clinical cancer research, a significant challenge is acquiring enough cellular material for direct drug screening, mainly due to the limited amount of initial tumour material available ^8,13^. Propagating cells through patient-derived xenografts has emerged as a particularly vital strategy for cultivating patient-derived cells from paediatric tumours ^25,68^, however this approach can take months and is often unsuccessful. Within a shorter 7–14-day period, we successfully expanded patient-derived cells in ECM-mimic hydrogels in a high-throughput fashion—an achievement not always possible with conventional tissue culture. Our workflow demonstrated significant potential for cell expansion and potential biobanking in the ECM-mimicking environments. We found that the necessary cell numbers can be attained using our hydrogel systems and high-throughput drug screening can be conducted directly *in situ*. Furthermore, bioprinting and cell expansion did not affect key molecular and phenotypic characteristics of the original tumours. Importantly, we were able to identify differential drug sensitivities in four out of seven samples all of which included one of more specific targeted agents.

Like the recently published HTP drug screening analyses of multiple paediatric patient cohorts, our study found that not all molecular vulnerabilities of patient-derived cells matched the specific drug susceptibilities identified in the ex vivo screens ^8,12,69^. This suggests that our HTP drug screening in the 3D bioprinted organoid models closely resembles pre-clinical drug testing outcomes and is in line with clinical observations that targeting a genetic alternation with a targeted drug does not necessarily result in an objective patient response ^10,11^. Our results confirm the importance of *ex vivo* HTP drug screening in support of clinical treatment decision making for paediatric patients and suggest that 3D bioprinted HTP drug screens could identify previously unrecognized chemotherapeutic and targeted drug vulnerabilities independent of tumour molecular profile.

Our study demonstrates the feasibility of applying this 3D bioprinting workflow to preclinical testing of patient-derived material. Additionally, this work contributes to ongoing efforts to develop patient organoid models and 3D technology for high-throughput drug screening, advancing their application into the preclinical setting. The significance of this work is that the 3D bioprinting workflow presented here can be extended to growing tumour organoids from a broad range of cancer types by incorporating elements of the native patient extracellular environment and could potentially be used for expansion of patient samples that are difficult to grow for subsequent biobanking. Furthermore, our approach could be of interest to a broader audience of developmental and cancer biologists where 3D cultures with defined physical, chemical, and biological characteristics are required to gain insights into cellular behaviours in response to various stimuli.

In conclusion, while the focus here was on high-risk neuroblastoma and sarcomas, preclinical model development and drug testing can be readily applied, adapted, and tested in the context of both paediatric and adult cancers. Our outlined workflow and platform can be adopted more broadly by researchers in the fields of cell and cancer biology. New insight into cellular growth requirements and impact on cell signalling and behaviour can be obtained, since the 3D bioprinting platform is compatible with a variety of downstream analyses including cell migration, invasion, and live cell imaging ^34^. By understanding and developing new ways to grow cells in an environment that mimics the patient tumour, we can improve predictability and identification of drug therapies for a broad range of cancer patients.

## Methods

### RESOURCE AVAILABILITY

#### Lead Contact

Further information, resources and reagent requests should be directed to and will be fulfilled by lead contact

#### Materials Availability

This study did not generate new unique reagents.

#### Data and Code Availability

This paper analyses existing patient WGS and RNAseq data which is available upon request from ZERO Childhood Cancer Program. Data access details are listed in the key resources table. Targeted sequencing data (TSO500), SNP and STR profile data, metabolic assay and microscopy data reported in this paper will be shared by the lead contact upon request. This paper does not report original code. Any additional information required to reanalyse the data reported in this paper is available from the lead contact upon request.

### EXPERIMENTAL MODEL AND STUDY PARTICIPANT DETAILS

#### Patient-Derived Samples

Patient-derived xenograft (PDX) models and patient DNA (zcc227) were obtained from ZERO Childhood Cancer Precision Medicine Program (ZERO). All procedures were approved by the Human Research Ethics Committee at the University of New South Wales (HC190693).

We selected a total of eight patient-derived samples with a range of patient and disease characteristics **(Table S1).**

#### Animal Studies

All animal studies and procedures were approved by the University of New South Wales Animal Care and Ethics Committee (ACEC 22/58B, ACEC 23/79B, ACEC 22/34B).

### METHOD DETAILS

#### Extracellular Matrix Gene Expression Analysis

RNA expression data from high-risk neuroblastoma (n=41); rhabdomyosarcoma (n=40); Ewing sarcoma (n=40) and osteosarcoma (n=25) patients were extracted from the ZERO dataset (accessed on 1st of March 2024). Low count genes (transcript count per million below 10 in half of the samples) were filtered out. ECM genes selected based on Extracellular Matrix Organization Gene Set (n=265) extracted using Harmonizome^36^. Extracellular matrix components were annotated with R (v4.2.3) and the MatrisomeAnalyzeR package^39^. 265 genes were ranked according to their TPM values and the top and bottom 30 genes were selected. Z-scores for the TPM values were calculated and a heatmap with hierarchical clustering was generated with the ComplexHeatmap^73^ and fastcluster^74^ packages in R. Further analysis was conducted on genes annotated as the core matrisome set (n=79) with the MatrisomeAnalyzeR package. A comparative gene expression analysis was performed for collagen 1 (COL1A1) and fibronectin (FN1). Welch’s t-test was used for statistical evaluation for both genes of interest, comparing these genes against the remaining core matrisome genes in the dataset.

#### Patient-Derived Xenograft Propagation

Briefly, 5–6-week-old female NOD.Cg-Prkdcscid Il2rgtm1Wjl/SzJAusb (NSG) mice were purchased from Australian BioResources (Moss Vale, NSW, Australia). The animals were housed in a pathogen-free environment in research animal cages with total volume of 400 cm² (Tecniplast, Italy) with air filters in positive pressure ventiracks. Environmental enrichment consisted of bedding, enviro-dry material, and igloos. Irradiated rat and mouse breeder cubes and water were supplied ad libitum. For inoculation with cryopreserved cells from previously dissociated patient-derived tumour xenografts, cells were suspended in serum-free RPMI medium (Life Technologies) and combined with growth factor-reduced Matrigel (Corning Inc., Corning, NY, USA) at a 1:1 volume ratio (1 million cells per mouse). Cells were subcutaneously implanted into the flanks of female NSG mice using a 27-gauge needle (Terumo, Tokyo, Japan). For tumour fragment implantation, mice were anesthetized, and a 5-mm horizontal incision was made on the dorsal surface, 10 mm anterior to the base of the tail, to create a subcutaneous pocket. A 2–5-mm tumour fragment, pre-soaked in growth factor-reduced Matrigel, was inserted into the pocket, and the incision was closed with a wound clip. Buprenorphine (0.1 mg/kg, intraperitoneal) (Provet, Castle Hill, Australia) was administered for analgesia. Tumour size was monitored twice weekly using Vernier callipers, calculated using the formula: Length×Width×Height/2. Mice were euthanized when the tumour size reached 1000 mm³ by CO_2_ overdose followed by cervical dislocation. Tumour tissues were then collected for tumour dissociation, bioprinting and histology studies.

#### Tumour Dissociation

Tumours were dissociated using tumour dissociation kits according to manufacturer’s instructions (Tumour Dissociation kit, human; Miltenyi Biotec, #130-095-929). Briefly, tumours were minced in RPMI media and transferred to gentleMACS tubes with enzymes A, H and R to be homogenized using gentleMACS Octo Dissociator. Cell suspension was strained through a 70 µm strainer and spun down at 1500 rpm for 5 minutes (acceleration/deceleration speed 6) before supernatant was removed. Red blood cells were lysed by resuspending the cell pellet in 1 x ACK Lysing buffer (155 mM Ammonium Chloride, 10 mM Potassium Bicarbonate, 0.1 mM EDTA) for 5 minutes, then spun down and supernatant discarded. Final pellet was resuspended in appropriate culture media for cell counting and cell suspension stored at 4°C while being used.

#### Mouse Cell Depletion

Mouse Cell depletion was conducted according to manufacturer’s instructions for Mouse Cell Depletion kit (Miltenyi Biotec, #130-104-694). Briefly, single cell tumour suspension was obtained by passing through a 70 µm strainer and pelleting cells. Per 2 million tumour cells, pellet was resuspended in 80 µL of 0.5% BSA in DPBS and mixed with 20 µL of Mouse Cell Depletion cocktail then incubated for 15 minutes at 4°C. Volume was adjusted to 500 µL using 0.5% BSA in DPBS and ran on autoMACS Pro Separator to magnetically separate positive fraction (mouse cells and dead cells) and negative fraction (human tumour cells).

#### Patient-Derived Cells

All bioprinting experiments were carried out using freshly isolated patient-derived cells obtained from tumour xenografts (see *patient-derived xenograft propagation* section above). Patient-derived neuroblastoma cells were cultured in vitro in IMDM media (Gibco, #12440061), 20% foetal bovine serum (FBS) and 1% Insulin-Transferrin-Selenium 100x (ITS-G, Gibco, #41400045). Sarcoma cells were cultured in alpha MEM (Gibco, #1251056) or IMDM media, 10% FBS, 1% ITS-G and 0.1% ROCK inhibitor (Y-27632 2HCl, SelleckChem, #S1049). All cells in vitro were cultured with 1% penicillin/streptomycin (Gibco) and 0.1% Amphotericin B (Gibco, #15290018).

#### Bioprinting

Bioinks and activators for multiple hydrogels were sourced from Inventia Life Science, Sydney, Australia (see table for further details). Hydrogels contain ECM-mimicking peptides of fibronectin (FN; RGD peptide), collagen 1 (CN; GFOGER peptide) or laminin (LN; DYIGSR peptide). 3D bioprinted models were printed using the RASTRUM 3D bioprinter (Inventia Life Science) as described in^33^. Printing protocols and hydrogel design were generated using RASTRUM Cloud (Inventia Life Science). For cell viability experiments, cells were primed (seeding density of 8,000 or 16,000 cells per well) before being printed in Imaging Models on flat bottom 96-well plates. Cells were bioprinted at these same priming densities using the Immunohistochemistry Models for immunohistochemistry experiments in standard 24-well plates. For sequencing experiments, cells were printed at either 36,000 or 72,000 cells per well in Large Plug Models in flat-bottom 96-well plates. For drug screening, cells were bioprinted in HTP Models in white clear bottom 384-well plates (Greiner, #781098) and printed at 2,000 or 4,000 cells per well.

| Hydrogel* | Stiffness | Mimicking component | Peptide/s | Catalogue Code |
| --- | --- | --- | --- | --- |
| 1.1kPa + FN | 1.1kPa | Fibronectin (FN) | RGD | Px02.31P |
| 1.1kPa + FN + CN | 1.1kPa | Fibronectin (FN)<br>Collagen 1 (CN) | RGD<br>GFOGER | Px02.09P |
| 1.1kPa + FN + CN+<br>LN | 1.1kPa | Fibronectin (FN)<br>Collagen 1 (CN)<br>Laminin (LN) | RGD<br>GFOGER<br>DYIGSR | Px02.28P |
| 3kPa + FN | 3kPa | Fibronectin (FN) | RGD | Px03.31P |
| 3kPa + FN + CN | 3kPa | Fibronectin (FN)<br>Collagen 1 (CN) | RGD<br>GFOGER | Px03.09P |
| 3kPa + FN + CN+<br>LN | 3kPa | Fibronectin (FN)<br>Collagen 1 (CN)<br>Laminin (LN) | RGD<br>GFOGER<br>DYIGSR | Px03.28P |

\*Hydrogels were obtained from Inventia Life Science and contain ECM-mimicking peptides of fibronectin (FN; RGD peptide), collagen 1 (CN; GFOGER peptide) or laminin (LN; DYIGSR peptide).

#### Cell Viability and Live/Dead Imaging

Tumour samples were bioprinted in each hydrogel condition using Imaging Models and cultured in 5% O_2_ 37°C/CO_2_ incubator for up to 14 days. Media was replenished at day 3, 7 and 10. Cells were incubated with Alamar Blue reagent (Sigma-Aldrich) at 10% media volume for 24 hours at day 1, 3, 7 and 14. The assay was read at 570-595nm using the Benchmark Plus plate reader (BIO-RAD) and absorbance values were normalized to Ethanol treated control wells (negative control). For Live/Dead imaging, cells were stained with Live/Dead viability/cytotoxicity kit, for mammalian cells (Invitrogen, #L3224) according to manufacturer’s instructions and previously published ^34^. Briefly, cells were bioprinted and cultured for up to 14 days. At day 7 and day 14, hydrogels were rinsed with DPBS and stained with 100 µL of live/dead staining solution (10 µM Ethidium Homodimer-1 and 5µM Calcein AM in DPBS) then incubated at 37°C for 30-45 minutes. Z-stack images were taken at 5x magnification using green fluorescence for live cells and red fluorescence for dead cells using CellDiscoverer 7 microscope (Zeiss). Images were visualized using Arivis 4D Vision software.

#### Cell Extraction

Cells were collected using the RASTRUM Cell Retrieval Solution, purchased from Inventia Life Science (#F235) according to manufacturer’s instructions. Cells were bioprinted in Px02.28P hydrogels and cultured until endpoint. Hydrogels were rinsed with DPBS and incubated with 50 µL Cell Retrieval Solution for 30 minutes at 37°C/5% CO_2_. Solutions were vigorously resuspended to dislodge cells from hydrogels and supernatant was collected in a 50 mL centrifuge tube through a 70 µm strainer to remove any hydrogel fragments. Wells were rinsed with DPBS and contents combined with the collected cell suspension. Cells were pelleted at 1200 rpm for 5 minutes then pellets rinsed twice with cold DPBS.

#### Sequencing and Sample Validation

##### Targeted Sequencing (TSO500)

Patient-derived cells were bioprinted and cultured for 7 or 14 days in relevant media, then cells were retrieved (see *Cell Extraction* section above) and pellets were snap frozen on dry ice and stored at −80°C for downstream analysis. Cell pellets were sent to Central Adelaide Local Health Network laboratories where DNA was extracted and sequenced using the Illumina TruSight Oncology 500 (TSO500) panel. DNA fastqs were first aligned to human and mouse genome simultaneously by bbsplit from BBTools (v38.79), outputting reads best mapping to each genome to two interleaved fastqs respectively. Human genome interleaved fastqs were converted to paired read fastqs by reformat from BBTools (v38.79) and used as input for TSO500 app. For TSO500 variant calling was performed using the Illumina TSO500 local app (v2.0). For other panels, variant calling was performed using Vardict (v1.8.2). After variant calling, variant annotation was performed using VEP (v100) and annotated with COSMIC (v95), ClinVar (2020-05-13), gnomAD (r2.1.1) and CADD (v1.6) using vcfanno. Potential pathogenic variants were identified using these databases and confirmed in OncoKB. CNV estimation was performed using CNVkit (v0.98) and tumour purity and ploidy were estimated using PureCN (v1.16) to turn fold change into estimated copy number. The pipeline was run on DNAnexus and genomic interpretation finalized using gentian, an in-house developed tool for assimilating and assessing targeted capture panel data.

##### STR and SNP validation

For samples requiring further validation, DNA was extracted from bioprinted cell pellets using the AllPrep DNA/RNA Micro Kit (QIAGEN, #80284) and DNA concentrations measured using a NanoDrop spectrophotometer. DNA precipitation was performed to further remove buffer salts by incubating sample DNA with 0.3 M Sodium Acetate and 2 volumes of 100% ethanol for 1 hour at −20°C. Samples were then rinsed in ice-cold 70% ethanol and resuspended in EB buffer. DNA samples were submitted to the Garvan Institute of Medical Research for short tandem repeat (STR) profiling using the PowerPlex® 18D system, which analyses 18 markers. The results were compared with STR profiles from the original tumour sample. Profiles with more than 80% identity were considered a match. Additionally, DNA samples were submitted to the Victorian Clinical Genetics Services for single nucleotide polymorphism (SNP) profiling using the Illumina Infinium Global Screening 531 Array-24 v2.0. The process involves two output metrics: Log R, indicating the overall signal intensity for both alleles, and B allele frequency (BAF), reflecting the allelic imbalance of single nucleotide polymorphisms (SNPs). To generate genome-wide allele-specific copy number profiles, the R package ASCAT (v2.5.2) was utilized for data analysis. Validation of the results was conducted by comparing the derived copy number profiles and purity with those of the original patient tumour samples.

##### PCR

Zcc227 patient-specific DNA sequence of EWSR-ERG breakpoint structural variant was obtained using WGS data. Custom oligonucleotide primers were designed using Integrated DNA Technologies PrimerQuest™ Tool (forward: GACTGCTAGCCCTGCTG; reverse: GTAAAGGAGCTGTGTTGCTTAT). Standard PCR was performed using AmpliTaq Gold™ DNA Polymerase from Thermo Fisher Scientific (#4311820) at a setting of 40 cycles and annealing temperature of 60^0^ C on the Bio-Rad T100 Thermo Cycler. PCR products were visualized against an in-house molecular ladder on an in-house 12.5% polyacrylamide gel using the Gel Doc XR+ System and Image Lab Software. PCR products were cleaned up with ExoSAP-IT™ (Thermo Fisher Scientific, #75001.1.ML).

##### In *Vivo* Tumourigenicity

To determine if the 3D bioprinted and expanded patient derived cells maintained their phenotypic characteristics to form tumours, we performed a tumourigenicity study as follows. Neuroblastoma zcc373 or Ewing Sarcoma zcc207 samples were propagated as described above and monitored until tumours reached endpoint at 1000 mm^3^. Tumours were collected, dissociated (see *Tumour Dissociation* section above) and bioprinted in Large Plug hydrogels at 36,000 cells per well. Remaining tumour cells were cryopreserved in 90% Fetal Bovine Serum (FBS) and 10% DMSO to be used as controls for tumourigenicity re-engraftment. Bioprinted samples were cultured for 10 days in relevant media before cells were extracted (see *Cell Extraction* section above). Freshly extracted bioprinted cells and thawed, matched cryopreserved PDX cells were resuspended in 100% Growth Factor Reduced Matrigel® (Corning, #354230) and subcutaneously inoculated at 1 million cells per mouse for each condition into the flanks of female (5-7 weeks old) NSG mice (NOD.Cg-Prkdcscid Il2rgtm1Wjl/SzJArc) obtained from Australian Resource Centre (Murdoch, Perth Western Australia). Once inoculated, tumour growth was monitored twice weekly using vernier calipers using the formula: ½ x L x W^2^. Mice were euthanized when the tumour size reached 1000 mm³ by CO_2_ overdose followed by cervical dislocation. Tumour tissues were then collected for histology and immunohistochemistry studies.

##### Immunohistochemistry

Patient-derived cells were bioprinted in Px02.28P hydrogels using the Immunohistochemistry Model and cultured for 14 days in relevant culture media. Bioprinted plates were fixed in 4% paraformaldehyde (PFA) in DPBS for 30 minutes at room temperature, then rinsed and stored in DPBS until use. For sample processing, hydrogels were first stained with 1:100 Trypan Blue in DPBS for 15 minutes. Structures were then carefully scooped out of plates and transferred to cryomold cassettes filled with O.C.T. compound (Tissue-Tek, #IA018) before being snap frozen on dry ice. Cryopreserved samples were processed by the Biological Specimen Preparation Laboratory in Katharina Gaus Light Microscopy Facility (KG-LMF) at the University of New South Wales into 5µM sections. Patient-derived xenograft tumours were collected at endpoint (1000 mm^3^) (see In *Vivo Tumourigenicity* section above), fixed in 10% formalin and paraffin embedded. Samples were cut into 5 µm sections. H&E and antibody marker staining was performed by the Histopathology department at the Garvan Institute of Medical Research using Leica Bond system. Tumour-specific markers were visualized with CD99 (BioCare Medical, #CME392A; 1:50), PHOX2B (abcam, #ab183741; 1:1000) and SATB2 (Cell Marque, #CM384R15; 1:100) antibodies. Ki-67 (SP6) (Thermo Fisher, #MA5-14520; 1:500) was used for cell proliferation.

##### High-Throughput Drug Screening

Patient-derived cells were bioprinted for drug screening (see *Bioprinting* section) and 50 µL of the relevant media added to each well. Prior to bioprinting, sarcoma samples were subject to mouse cell depletion protocol (see *Mouse Cell Depletion* section). All samples were printed in two identical plates (duplicates). Media was exchanged and replaced with 50 µL media every 3-4 days for duration of experiment. After 7 or 14 days of incubation, a library of 48 compounds (Compounds Australia) was administered to duplicate HTP plates using the Hamilton STAR liquid handling robot in final concentrations ranging from 0.5 nmol/L-5 µmol/L. Drug compounds were selected based on previously identified differential drug sensitivities in specific patient samples^13,15^, standard of care chemotherapy agents against neuroblastoma and sarcoma tumours and included previously unconsidered targets of potential interest. Benzethonium chloride (100 µmol/L) and DMSO were administered as positive and negative controls, respectively. 72 hours after drug administration, metabolic activity was measured using the CellTiter-Glo® 3D Cell Viability Assay (Promega, #G9683). Briefly, 50 µL of CellTiter-Glo® reagent was added to each well (1:1 to media volume) and shaken on plate shaker at 450 rpm for 2 hours, protected from light. Luminescence was read on the EnVision® plate reader (PerkinElmer) for 1 second per well. Data analysis was performed as previously described^13^. ActivityBase (IDBS) software (Version 8.3.0.175) was used to generate cell viability percentages from raw luminescence data using the following formula: ([readout value drug – average readout of positive controls] / [average readout of negative ^controls^ – average readout of positive controls]) × 100. Calculated viabilities were used to generate dose-response curves, calculate area under the curve (AUC) and half-maximal inhibitory concentration (IC50) values.

### QUANTIFICATION AND STATISTICAL ANALYSIS

Extracellular matrix gene expression analysis was performed using R (v4.2.3) and the MatrisomeAnalyseR package. Welch’s t-test was used for statistical evaluation of gene expression compared against remaining core matrisome genes in the dataset. Cell viability data is presented as mean +/- SD, and representative images shown from one of three independent experiments. Sample size (n) values are provided in associated main and supplementary figures. Statistical analyses were performed in GraphPad Prism software (v.10.3.0).

## Acknowledgments

Children’s Cancer Institute Australia is affiliated with University of New South Wales and the Sydney Children’s Hospital Network. We acknowledge ZERO Childhood Cancer personalized medicine program for providing samples and molecular data related to this project. The authors would like to thank all the patients, parents and healthcare professionals that participated in ZERO Childhood Cancer program. We would like to thank Biljana Dumevska from ZERO Core Management for her assistance in coordinating data and samples access. The authors thank the Children’s Cancer Institute Animal Facility for providing support to this study

This work is supported by a National Health and Medical Research Council (NHMRC) Investigator Grants (#2016464 to M.K, #1196648 to J.J.G.), NHMRC Synergy Grant #2019056 to M.K and J.J.G. Cancer Council New South Wales #2020797, and Cancer Australia #1185313 led by M.K. We thank the Luminesce Alliance for supporting the bioinformatics analysis.

We are greatly appreciative of our consumer representatives Amanda Younes, Josiane Demetriou, Darrin Batchelor and Simon Sleep for their support and providing valuable feedback on the content and consumer relevance of this research. We thank Julio Ribeiro, Robert Utama, Aidan O’Mahony and Inventia Life Science team for their technical support and advice on the 3D bioprinting. We would like to acknowledge Fei Shang and Maria Kasherman from Katharina Gaus Light Microscopy Unit at the Mark Wainwright Analytical Centre at UNSW Sydney for assistance with hydrogel cryosectioning and imaging. Anaiis Zaratzian, Andrew M. Da Silva, Michael Tayao from Garvan Institute Histopathology Facility for the histology services. Madeleine Wheatley, Joan Solomon and Brandon Hearn for technical assistance with animal experiments. Roxy Cadiz and Karina Pazaky for technical assistance with high-throughput drug screening. Michelle Henderson, Libby Huang, Luis Enriquez and Ani Lack for technical advice and assistance with PCR validation experiments. We would like to thank Claudia Flemming and Omesha Perera for their assistance in the manuscript preparation. Schematics for methods and results were created with BioRender.com.

## Author Contributions

Conceptualization, M.K.; and M.J.; Methodology, M.J., J.N.S., S.G., G.T., A.K., A.J.X., K.K., C.M., E.D.G.F., A.M.F., Z.B., M.J.K, J.I.F., M.E.M.D., J.J.G. and M.K.; Software, P.G., J.M., M.W.E., L.C., P.V., C.M., J.G. and E.Y.D., M.J.K; Formal Analysis, V.P., M.J., J.N.S., G.T., P.G., J.M., M.W.E., L.C., K.K., P.V., M.J.K., M.E.M.D. and M.K.; Investigation, V.P., M.J., J.N.S., S.G., A.K., K.K., L.C. and A.J.G.; Resources, A.K., A.J.X., L.C., M.E.M.D.; Writing – Original Draft, V.P., M.J., J.N.S, M.E.M.D and M.K.; Writing – Review & Editing, V.P., M.J., J.N.S., A.J.G., L.M.S.L., M.E.M.D., J.J.G. and M.K.; Visualization, V.P., M.J., J.N.S., P.G.,; Supervision, M.K.; Project Administration, V.P., M.J., J.N.S. and M.K.; Funding Acquisition, M.K.

## The Paper Explained

### Problem

Children with high-risk cancers have limited treatment options. Personalised drug treatments, tailored to individual genetic profiles, are becoming more popular. However, paediatric tumour biopsies often don’t provide enough material beyond standard diagnostics. This lack of material makes it difficult to conduct comprehensive personalised drug testing, which requires cell expansion through primary culture or patient-derived xenograft models (PDXs). These methods often take a long time and have limited success. Additionally, there are no high-throughput paediatric tumour models that replicate the tumour’s natural environment while allowing for timely, high-throughput direct drug screening for paediatric patients.

### Results

We introduce a proof-of-concept using high-throughput 3D bioprinting and patient-derived cancer cells. We developed a platform that expands freshly isolated tumour cells in engineered hydrogels that mimic the extracellular matrix (ECM) of the tumour environment. Using ECM-mimic hydrogels, we created patient-specific tumour organoids in their ‘native’ growth conditions. These organoids retained the genetic and phenotypic traits of the original tumours and remained tumourigenic. Direct drug screening in bioprinted organoids identified individualised sensitivities, offering a timely, clinically relevant platform for precision medicine in paediatric cancers.

### Impact

Our platform enables robust, high-throughput expansion of freshly isolated tumour cells in engineered ECM-mimic hydrogels for personalised medicine. The tumor organoids provide a powerful tool for individualised drug sensitivity profiling, with the potential to transform paediatric preclinical testing and expand therapeutic options for patients in a clinically relevant timeframe. Our platform offers an innovative approach for expanding patient-derived cells for direct preclinical drug testing in paediatric cancers and could be applied to advance precision medicine practices and biological discovery across various cancer types.

## Supplemental information

**Supplementary Figure 1 Cell proliferation of 3D bioprinted patient-derived cells.** Cell proliferation graphs of 3D bioprinted patient-derived cells in the hydrogel conditions. Each sample was bioprinted in either 1.1kPa or 3kPa hydrogels, containing fibronectin (FN) only or FN and collagen (CN) peptides. Proliferation rate was measured at Day 1, 3, 7 and 14 post-printing. All experiments were repeated three times. Graphs are separated into cancer types A) neuroblastoma, B) Ewing sarcoma and C) osteosarcoma. Data are representative of mean ±SD. Related to Figure 1.

**Supplementary Figure 2 Cell viability of 3D bioprinted patient-derived cells.** Each sample was bioprinted in 3kPa + FN + CN + LN hydrogels and cultured up to 14 days. Cells were stained with calcein-AM (green; live)/ethidium homodimer 1 (red; dead) Live/Dead Assay. Z stack 3D images were taken at day 14 post-printing. Representative image shown for each patient sample in brightfield (left) and Live/Dead (right). Scale bars on all images are 500 µm. Related to Figure 1.

**Supplementary Figure 3** 3D bioprinted tumour organoids are compatible with a high-throughput preclinical drug screening platform and reveal patient-specific drug vulnerabilities. Log2[IC50] distribution across all samples for a 48-drug library. Drugs were ordered based on the lowest to highest median Log2[IC50] values, followed by lowest quartile and then lowest detected Log2[IC50] values. Related to Figure 6.

