## Supplementary figures and images for "Engineered paediatric tumours retain maintains tumour genotype and phenotype for precision medicine"

### Suppl Figs 1-3

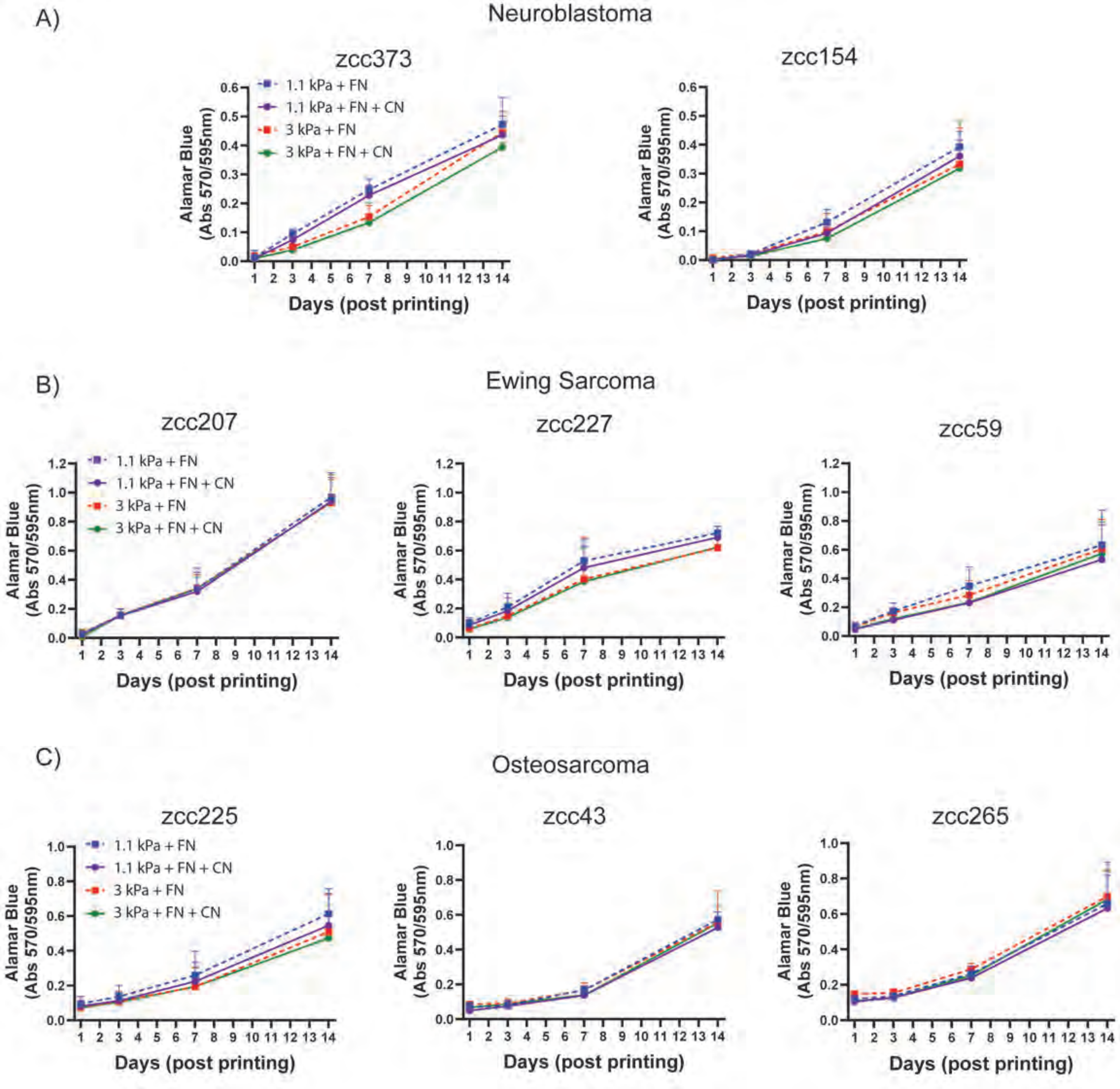

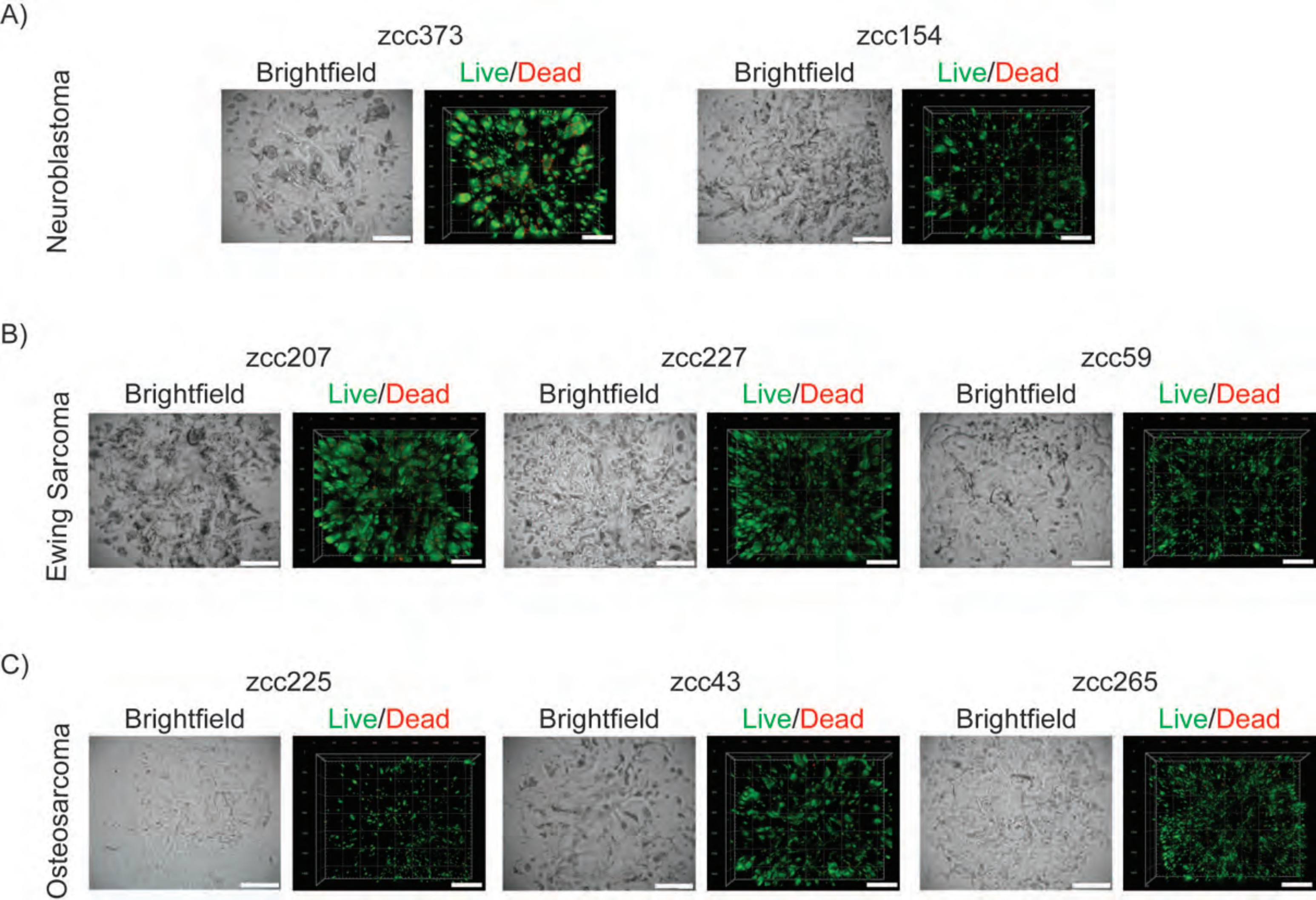

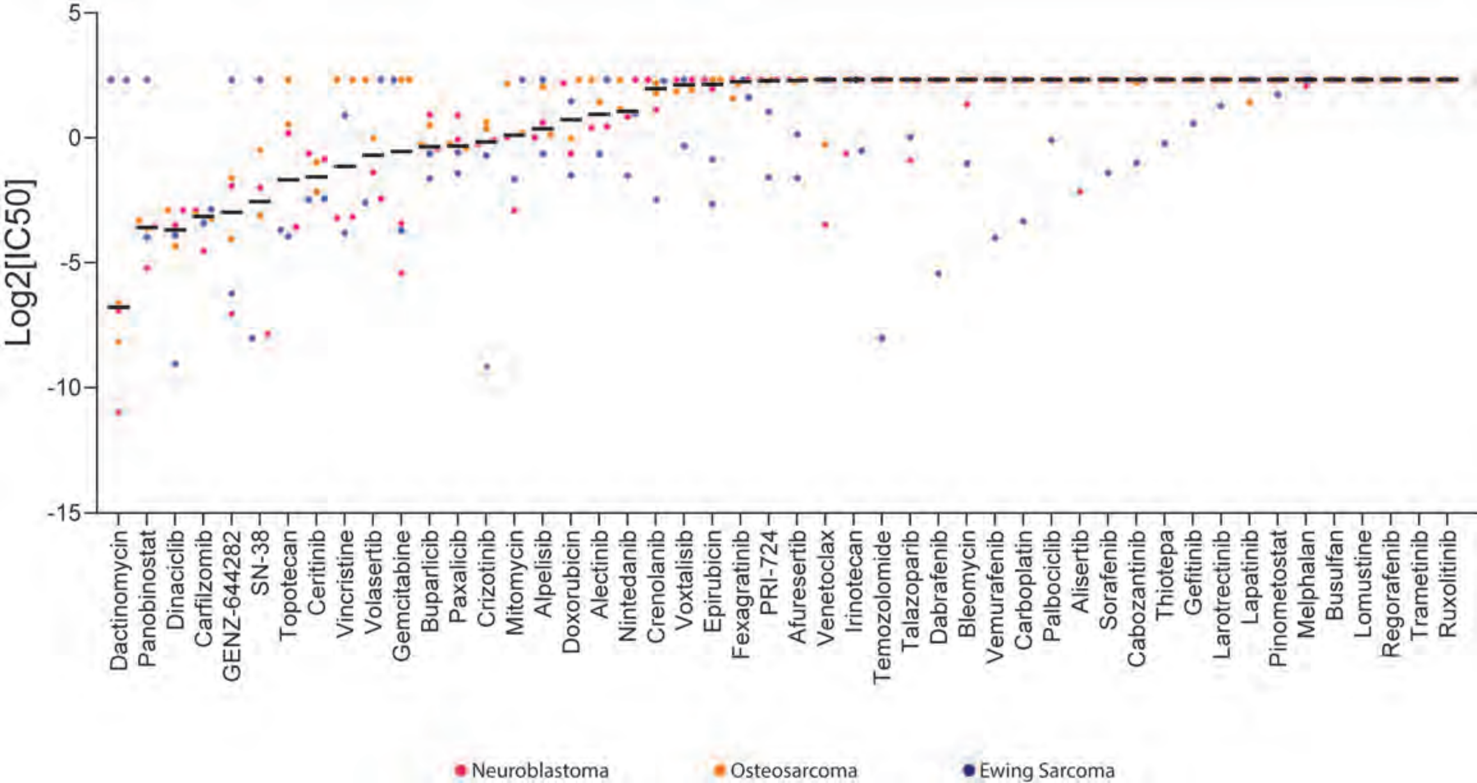
