## Supplementary material for "Engineered paediatric tumours retain maintains tumour genotype and phenotype for precision medicine": Suppl Tables 1-6

Supplementary Table 1 Patient demographics and patient-derived sample information, related to Figure 2.

| Cancer Type | Sample ID | Sex/Age (years) | Sample Type | Key molecular characteristics |
|---|---|---|---|---|
| **Neuroblastoma** | zcc154 | M/7 | Progression | *MYCN* amplification; *ALK* focal amplification |
| | zcc373 | M/1 | Relapse | *MYCN* amplification |
| **Ewing Sarcoma** | zcc59 | F/12 | Relapse | Somatic *PIK3CA* mutation; *EWSR1-ETV1* fusion |
| | zcc207 | F/14 | Diagnostic | Somatic *TP53* mutation; *TP53* gain; *EWSR1-FL1* fusion |
| | zcc227 | M/5 | Relapse | *EWSR1 – ERG* fusion |
| **Osteosacoma** | zcc43 | F/15 | Diagnostic | *TP53 - LSMD1* duplication |
| | zcc225 | F/17 | Relapse | *NUDT2 -TP53* fusion; *DLG2-DLG2* deletion |
| | zcc265 | F/18 | Relapse | *RB1* germline mutation; *DLG2- DLG2* segmental deletion; *TP53-TP53* segmental biallelic deletion |

### Supplementary Table 2 Success rates for generating cultures from patient-derived cells (fresh or cryopreserved)

| Cancer Type | Attempted cultures for unique ZERO* samples | Samples with successful cultures | Succes (%) |
|---|---|---|---|
| **Neuroblastoma** | 10 | 0 | 0 |
| **Ewing Sarocoma** | 21 | 1 | 4.8 |
| **Osteosacoma** | 23 | 3 | 13 |

* ZERO Childhood Cancer Precision Medicine Program

Supplementary Table 3 In vivo expansion time for individual patient-derived samples, related to Figure 2.

| Cancer Type | Sample ID (ZERO*) | Time (days**) |
|---|---|---|
| **Neuroblastoma** | zcc154 | >70 |
| | zcc373 | >88 |
| **Ewing Sarcoma** | zcc59 | >57 |
| **Osteosacoma** | zcc43 | >98 |
| | zcc265 | >179 |

* ZERO Childhood Cancer Precision Medicine Program
**In vivo expansion time was only available for five out of eight patient samples

Supplementary Table 4 Comparative analysis of genetic variants in original patient tumours by whole genome sequencing and 3D tumouroids by targeted sequencing*

| Disease | Patient | Gene | Variation | Description | Status | Original patient sample | | | Bioprinted sample | | |
|---|---|---|---|---|---|---|---|---|---|---|---|
| | | | | | | Whole Genome Sequencing (WGS) | Copy number | Variant allele frequency (VAF) | TruSight500 (TSO500) | Copy number | Variant allele frequency (VAF) |
| Neuroblastoma | zcc373 | *MYCN* | Somatic copy number variant | CNV | pathogenic | Yes | 201.51 | | Yes | 73.6 | |
| | zcc154 | *MYCN* | Somatic copy number variant | CNV | pathogenic | Yes | 49.82 | | Yes | 127.5 | |
| | | *ALK* | Somatic copy number variant | CNV | pathogenic | Yes | 66.34 | | Yes | 80.6 | |
| | | *ALK-THADA* | Somatic structural variant (duplication) | SV | not pathogenic | Yes | | | Yes | | |
| Ewing Sarcoma | zcc207 | *EWSR1-FLI1* | Somatic structural variant (fusion) | SV | pathogenic | Yes | | | Yes | | |
| | | *STAG2* | Single nucleotide variation NM_006603.4(STAG2):c.3395T>G (p.Leu1132Ter) | SNV | likely pathogenic | Yes | | 43.24% | Yes | | 63.10% |
| | | *TERT* | Single nucleotide variation NM_198253.2(TERT):c.-57A>C | SNV | likely pathogenic | Yes | | 65.09% | Yes | | 32.35% |
| | | *TP53* | Single nucleotide variation NM_000546.5(TP53):c.577C>T (p.His193Tyr) | SNV | pathogenic | Yes | | 85.37% | Yes | | 99.84% |
| | zcc227 | *EWSR-ERG* | Somatic structural variant (fusion) | SV | pathogenic | Yes | | | No | | |
| | zcc59 | *SMARCA4* | Single nucleotide variation NM_003072(SMARCA4):c.3469C>T (p.Arg1157Trp) | SNV | likely pathogenic | Yes | | 4.58% | Yes | | 47.09% |

| | | | | | | | | | | | |
|---|---|---|---|---|---|---|---|---|---|---|---|
| | | *PIK3CA* | Single nucleotide variation NM_006218.4(PIK3CA):c.3140A>G (p.His1047Arg) | SNV | pathogenic | Yes | | 8.09% | Yes | | 36.40% |
| | | *ARID1A* | Single nucleotide variation NM_006015(ARID1A):c.3859dupA (p.Arg1287LysfsTer11) | SNV | pathogenic | Yes | | 21.21% | Yes | | 18% |
| | | *EWSR1 - ETV1* | Somatic structural variant (fusion) | SV | pathogenic | Yes | | | Yes | | |
| **Osteosarcoma** | **zcc225** | *NUDT21-TP53* | Somatic structural variant (fusion) | SV | reportable | Yes | | | Yes | | |
| | **zcc43** | *TP53-LSMD1* | Somatic structural variant (fusion) | SV | reportable | Yes | | | No | | |
| | **zcc265** | *BCL2* | Somatic copy number variant | CNV | pathogenic | Yes | 17.59 | | Yes | 17.1 | |
| | | *RB1* | Single nucleotide variation NM_000321(RB1):c.507_508dupTG (p.Glu170ValfsTer6) | SNV | pathogenic | Yes | | 99.83% | Yes | | 97% |
| | | *TP53-TP53* | Structural variant<br>Segmental biallelic deletion (exon 1) | SV | pathogenic | Yes | | | No | | |

* TruSight Oncology 500 (TSO500)

Table S5. Drug library information, related to Figure 6.

| Drug | Synonyms | Category | Drug Class | Target(s) |
|---|---|---|---|---|
| Afuresertib | GSK-2110183; GSK2110183 | Targeted | PI3K-AKT-mTOR signaling | AKT1; AKT2; AKT3 |
| Alectinib | Alecensa; RG7853; AF802; CH5424802; CH 5424802; RO5424802 | Targeted | Receptor tyrosine kinase signaling | ALK |
| Alisertib | MLN8237 | Targeted | Aurora-PLK signaling | AURKA |
| Alpelisib | BYL-719; BYL719 | Targeted | PI3K-AKT-mTOR signaling | PI3Ka |
| Bleomycin | Bleomycin sulfate; Blexane; NSC125066; Blenoxane | Chemotherapeutic | DNA structure and function | Free radical-promoting agent |
| Buparlisib | BKM120; NVP-BKM120 | Targeted | PI3K-AKT-mTOR signaling | PI3Ka; PI3Kb; PI3Kd; PI3Kg |
| Busulfan | Myleran; Busulphan; Sulphabutin; Myelosan; Leucosulfan; Busulfex | Chemotherapeutic | DNA synthesis | DNA dialkylating agent (N7-guanine residues) |
| Cabozantinib | XL184; XL-184; BMS-907351; Cometriq | Targeted | Receptor tyrosine kinase signaling | c-Met; VEGFR2; AXL; RET; c-Kit; Flt3 |
| Carboplatin | NSC 241240 | Chemotherapeutic | DNA structure and function | DNA crosslinker |
| Carfilzomib | PR-171 | Targeted | Proteasome function | Proteasome |
| Ceritinib | LDK378 | Targeted | Receptor tyrosine kinase signaling | ALK |
| Crenolanib | CP-868596 | Targeted | Receptor tyrosine kinase signaling | PDGFRa; PDGFRb; Flt3 |
| Crizotinib | PF-02341066; PF02341066; PF 02341066; Xalkori | Targeted | Receptor tyrosine kinase signaling | c-Met; ALK; ROS1 |
| Dabrafenib | GSK2118436 | Targeted | MAPK-ERK signaling | BRAFV600E |
| Dactinomycin | Actinomycin D | Chemotherapeutic | RNA synthesis | dsDNA intercalator |
| Dinaciclib | SCH727965; SCH-727965; SCH 727965; MK-7965; PS-095760 | Targeted | Cell cycle-checkpoint signaling | CDK1; CDK2; CDK5; CDK9 |
| Doxorubicin | Doxorubicin hydrochloride; Adriamycin; Adriacin; Adriblastina; Adriblastin | Chemotherapeutic | DNA replication | DNA topoisomerase II inhibitor |
| Epirubicin | Epirubicin hydrochloride; 4'-epidoxorubicin hydrochloride | Chemotherapeutic | DNA replication | DNA topoisomerase II inhibitor |
| Fexagratinib | AZD4547 | Targeted | Receptor tyrosine kinase signaling | FGFR1; FGFR2; FGFR3; FGFR4; KDR |
| Gefitinib | ZD1839; ZD-1839 | Targeted | Receptor tyrosine kinase signaling | EGFR |
| Gemcitabine | Gemcitabine hydrochloride; Gemzar; LY-188011 | Chemotherapeutic | Nucleic acid synthesis or utilization | Antimetabolite (pyrimidine analogue) |
| GENZ-644282 | genz644282; Genz 644282 | Chemotherapeutic | DNA replication | DNA topoisomerase I inhibitor |
| Irinotecan | Irinotecan hydrochloride; (+)-Irinotecan; CPT-11 | Chemotherapeutic | DNA replication | DNA topoisomerase I inhibitor |
| Lapatinib | Tyverb; GW-572016, GSK572016; | Targeted | Receptor tyrosine kinase signaling | EGFR; HER2 |
| Larotrectinib | Larotrectinib sulfate; LOXO-101; ARRY-470 | Targeted | Receptor tyrosine kinase signaling | TrkA; TrkB; TrkC |
| Lomustine | CCNU | Chemotherapeutic | DNA structure and function | DNA alkylating agent (O6/N7-guanine residues) |
| Melphalan | Melphalan hydrochloride; Alkeran; Sarcolysin; L-PAM | Chemotherapeutic | DNA structure and function | DNA alkylating agent (N7-guanine residues) |
| Mitomycin C | Ametycine | Chemotherapeutic | DNA structure and function | DNA alkylating agent (N2-guanine residues) |
| Nintedanib | BIBF 1120; Intedanib | Targeted | Receptor tyrosine kinase signaling | VEGFR1; VEGFR2; VEGFR3; FGFR1; FGFR2; FGFR3; PDGFRa; PDGFRb |
| Palbociclib | PD0332991; PD-0332991; PF-00080665-73 | Targeted | Cell cycle-checkpoint signaling | CDK4; CDK6 |
| Panobinostat | LBH589 | Targeted | Epigenetic regulation | pan-HDAC |
| Paxalisib | GDC-0084; RG7666 | Targeted | PI3K-AKT-mTOR signaling | PI3Ka; PI3Kb; PI3Kd; PI3Kg; mTOR |
| Pinometostat | EPZ-5676; EPZ5676; EPZ 5676 | Targeted | Epigenetic regulation | DOT1L |
| PRI-724 | | Targeted | Wnt signaling | bCatenin-CBP |
| Regorafenib | BAY 73-4506; Stivarga | Targeted | Multitargeted | VEGFR1; VEGFR2; VEGFR3; PDGFRb; c-Kit; RET; Raf-1 |
| Ruxolitinib | INCB018424; INCB-18424; Jakafi | Targeted | JAK-STAT signaling | JAK1; JAK2 |
| SN-38 | | Chemotherapeutic | DNA replication | DNA topoisomerase I inhibitor |
| Sorafenib | BAY 43-9006; Nexavar; 284461-73-0 | Targeted | Multitargeted | Raf-1; B-Raf; VEGFR-2 |
| Talazoparib | BMN 673; BMN-673; LT-673 | Targeted | DNA damage signaling | PARP1; PARP2 |
| Temozolomide | TMZ; Temodar; Temodal; Temcad; SCH 52365 | Chemotherapeutic | DNA structure and function | DNA alkylating agent (O6/N7-guanine residues) |
| Thiotepa | Thio-TEPA; thioplex; tiofosfamid | Chemotherapeutic | DNA structure and function | DNA alkylating agent (N7-guanine residues) |
| Topotecan | Topotecan hydrochloride; K&F 104864-A; SKF 104864A; SKFS 104864A; NSC 609669; nogitecan hydro | Chemotherapeutic | DNA replication | DNA topoisomerase I inhibitor |
| Trametinib | GSK-1120212; GSK1120212; JTP-74057; Mekinist | Targeted | MAPK-ERK signaling | MEK1; MEK2 |
| Vemurafenib | RG7204; R7204; RO5185426; PLX4032 | Targeted | MAPK-ERK signaling | BRAFV600E |
| Venetoclax | ABT199; ABT-199; GDC-0199 | Targeted | Apoptotic signaling | BCL-2 |
| Vincristine | Vincristine sulfate; Leurocristine sulfate; 22-oxovincaleukoblastine sulfate | Chemotherapeutic | Mitosis | Microtubule destabilizer |
| Volasertib | BI6727; BI 6727; BI-6727 | Targeted | Aurora-PLK signaling | PLK1 |
| Voxtalisib | SAR 245409; SAR245409; XL765; XL-765 | Targeted | PI3K-AKT-mTOR signaling | PI3Ka; PI3Kb; PI3Kd; PI3Kg; DNA-PK; mTORC1; mTORC2 |

Table S6. AUC and Log2[IC50] T-score values for an individual drug and sample, related to Figure 6.

| | AUC T-score | | | | | | log2[IC50] T-score | | | | | |
|---|---|---|---|---|---|---|---|---|---|---|---|---|
| | NB | NB | EWS | EWS | OST | OST | NB | NB | EWS | EWS | OST | OST |
| **Drug** | **zcc154** | **zcc373** | **zcc207** | **zcc227** | **zcc225** | **zcc265** | **zcc154** | **zcc373** | **zcc207** | **zcc227** | **zcc225** | **zcc265** |
| Afuresertib | -0.07 | 1.40 | -0.85 | -1.18 | 0.93 | -0.23 | 0.62 | 0.62 | -0.69 | -1.74 | 0.62 | 0.58 |
| Alectinib | -0.95 | 0.87 | -1.39 | 0.29 | 1.15 | 0.03 | -0.49 | -0.54 | -1.44 | 1.07 | 1.07 | 0.32 |
| Alisertib | -0.03 | -1.86 | -0.07 | 0.24 | 0.86 | 0.87 | 0.41 | -2.04 | 0.41 | 0.41 | 0.41 | 0.41 |
| Alpelisib | -0.36 | -0.52 | -1.31 | 1.66 | 0.33 | 0.19 | -0.11 | -0.62 | -1.16 | 1.33 | 1.10 | -0.53 |
| Bleomycin sulfate | 0.05 | 0.67 | -1.04 | -1.32 | 0.40 | 1.25 | -0.20 | 0.54 | 0.54 | -1.95 | 0.54 | 0.54 |
| Buparlisib (BKM120) | 0.76 | -0.21 | 0.02 | -1.85 | 0.37 | 0.90 | 1.32 | -0.19 | -0.42 | -1.53 | -0.01 | 0.84 |
| Busulfan | 0.19 | 1.03 | -0.28 | -1.28 | -0.87 | 1.20 | 0.00 | 0.00 | 0.00 | 0.00 | 0.00 | 0.00 |
| Cabozantinib | -1.17 | 1.06 | -0.10 | -1.21 | 0.85 | 0.57 | 0.42 | 0.42 | 0.42 | -2.04 | 0.42 | 0.34 |
| Carboplatin | -0.31 | 0.70 | 0.18 | -1.88 | 0.62 | 0.70 | 0.41 | 0.41 | 0.41 | -2.04 | 0.41 | 0.41 |
| Carfilzomib | -1.93 | 0.35 | -0.07 | 0.88 | 0.19 | 0.58 | -1.93 | 0.68 | -0.13 | 0.77 | 0.44 | 0.17 |
| Ceritinib | 1.52 | 0.78 | -0.89 | -0.87 | 0.16 | -0.70 | 0.86 | 1.12 | -1.00 | -1.02 | 0.71 | -0.67 |
| Crenolanib | 0.86 | 0.08 | 0.51 | -1.96 | 0.14 | 0.37 | 0.60 | -0.04 | 0.58 | -1.98 | 0.31 | 0.53 |
| Crizotinib | 0.60 | 0.48 | -0.04 | -1.99 | 0.54 | 0.40 | 0.38 | 0.35 | 0.22 | -2.03 | 0.57 | 0.51 |
| Dabrafenib | -0.06 | 0.68 | -0.10 | -1.89 | 0.68 | 0.68 | 0.41 | 0.41 | 0.41 | -2.04 | 0.41 | 0.41 |
| Dactinomycin | -0.13 | -0.98 | -0.49 | 1.93 | -0.09 | -0.25 | -0.40 | -1.12 | 1.24 | 1.24 | -0.35 | -0.62 |
| Dinaciclib | 0.17 | 0.97 | 0.11 | -1.79 | -0.30 | 0.85 | 0.40 | 0.65 | 0.23 | -1.98 | 0.05 | 0.65 |
| Doxorubicin hydrochloride | -0.36 | 0.19 | -1.49 | -0.41 | 0.66 | 1.42 | 0.82 | -0.80 | -1.31 | 0.40 | NA | 0.90 |
| Epirubicin hydrochloride | -0.12 | 0.61 | -1.55 | -0.75 | 0.91 | 0.90 | 0.66 | 0.49 | -0.82 | -1.66 | 0.66 | 0.66 |
| Fexagratinib (AZD4547) | 0.86 | -0.42 | -0.84 | 1.61 | -0.75 | -0.47 | 0.75 | 0.75 | -1.23 | 0.75 | -1.31 | 0.30 |
| Gefitinib | -0.23 | 1.24 | -0.19 | -0.37 | -1.47 | 1.02 | 0.41 | 0.41 | 0.41 | -2.04 | 0.41 | 0.41 |
| Gemcitabine hydrochloride | -0.86 | -1.20 | -0.63 | 0.76 | 0.86 | 1.06 | -0.69 | -1.24 | -0.76 | 0.90 | 0.90 | 0.90 |
| GENZ-644282 | 0.27 | -1.14 | -0.91 | 1.61 | 0.38 | -0.21 | 0.34 | -1.15 | -0.91 | 1.57 | 0.43 | -0.28 |
| Irinotecan hydrochloride | 0.22 | -1.40 | -1.10 | 0.91 | 0.60 | 0.76 | 0.65 | -1.32 | -1.26 | 0.65 | 0.65 | 0.65 |
| Lapatinib | -0.18 | 1.31 | -1.52 | 0.57 | 0.46 | -0.64 | 0.41 | 0.41 | 0.41 | 0.41 | 0.41 | -2.04 |
| Larotrectinib sulfate | -0.76 | 0.33 | -0.90 | -0.52 | 0.05 | 1.80 | 0.41 | 0.41 | 0.41 | -2.04 | 0.41 | 0.41 |
| Lomustine | 0.88 | -0.34 | 0.63 | -1.20 | -1.05 | 1.09 | 0.00 | 0.00 | 0.00 | 0.00 | 0.00 | 0.00 |
| Melphalan hydrochloride | -1.15 | 0.31 | -1.36 | 0.52 | 0.69 | 0.99 | 0.41 | -2.04 | 0.41 | 0.41 | 0.41 | 0.41 |
| Mitomycin C | 0.28 | -1.50 | -0.93 | 0.37 | 0.65 | 1.13 | 0.00 | -1.42 | -0.80 | 1.12 | 0.08 | 1.04 |
| Nintedanib | 0.64 | 0.64 | -0.13 | -1.88 | 0.78 | -0.06 | -0.12 | 0.93 | -0.02 | -1.80 | 0.93 | 0.09 |
| Palbociclib | -0.10 | -0.26 | 0.31 | -1.36 | NA | 1.40 | 0.45 | 0.45 | 0.45 | -1.79 | NA | 0.45 |
| Panobinostat | -0.11 | -0.92 | -0.78 | 1.86 | 0.14 | -0.19 | -0.25 | -0.88 | -0.41 | 1.97 | -0.16 | -0.28 |
| Paxalisib (GDC-0084) | -0.98 | 1.25 | -1.41 | 0.19 | 0.38 | 0.57 | 0.31 | 1.60 | -1.48 | -0.37 | -0.14 | 0.07 |
| Pinometostat | -1.47 | 1.05 | 0.24 | -0.47 | -0.47 | 1.11 | 0.41 | 0.41 | 0.41 | -2.04 | 0.41 | 0.41 |
| PRI-724 | -0.03 | 0.99 | -0.07 | -1.88 | 0.51 | 0.48 | 0.56 | 0.56 | -0.25 | -1.93 | 0.56 | 0.51 |
| Regorafenib | 0.50 | -0.82 | -1.00 | 1.08 | 1.08 | -0.84 | 0.00 | 0.00 | 0.00 | 0.00 | 0.00 | 0.00 |
| Ruxolitinib (INCB018424) | 0.40 | 0.47 | -1.56 | -0.91 | 0.61 | 1.00 | 0.00 | 0.00 | 0.00 | 0.00 | 0.00 | 0.00 |
| SN-38 | -0.04 | -1.14 | -1.18 | 1.30 | 0.67 | 0.39 | 0.29 | -1.14 | -1.18 | 1.34 | 0.66 | 0.02 |
| Sorafenib | 0.59 | -0.14 | 0.54 | -1.96 | 0.43 | 0.55 | 0.41 | 0.41 | 0.41 | -2.04 | 0.41 | 0.41 |
| Talazoparib | -0.52 | -0.98 | -1.13 | 1.21 | 0.87 | 0.54 | 0.63 | -1.59 | -0.94 | 0.63 | 0.63 | 0.63 |
| Temozolomide | 0.46 | 0.45 | 0.42 | -2.04 | 0.44 | 0.27 | 0.41 | 0.41 | 0.41 | -2.04 | 0.41 | 0.41 |
| Thiotepa | 1.02 | 0.37 | -0.73 | -1.48 | -0.21 | 1.02 | 0.41 | 0.41 | 0.41 | -2.04 | 0.41 | 0.41 |
| Topotecan hydrochloride | -0.41 | -0.52 | -0.62 | -0.97 | 1.09 | 1.43 | 0.57 | -0.81 | -0.86 | -0.96 | 1.36 | 0.70 |
| Trametinib | -1.85 | 0.42 | -0.32 | 0.73 | 0.18 | 0.85 | 0.00 | 0.00 | 0.00 | 0.00 | 0.00 | 0.00 |
| Vemurafenib | 0.56 | 0.40 | 0.30 | -2.03 | 0.43 | 0.35 | 0.41 | 0.41 | 0.41 | -2.04 | 0.41 | 0.41 |
| Venetoclax | -0.28 | -1.75 | 0.01 | 0.89 | 0.11 | 1.02 | 0.58 | -1.84 | 0.58 | 0.58 | -0.50 | 0.58 |
| Vincristine sulfate | -0.92 | -0.39 | -1.31 | 0.88 | 0.91 | 0.84 | -0.83 | -0.82 | -1.04 | 0.57 | 1.06 | 1.06 |
| Volasertib | -0.57 | -0.83 | -1.25 | 0.93 | 1.00 | 0.73 | -0.49 | -0.96 | -1.03 | 1.17 | 1.17 | 0.13 |
| Voxtalisib | 1.20 | 0.01 | -0.33 | -1.72 | 0.16 | 0.68 | 0.57 | 0.57 | -2.00 | 0.57 | 0.12 | 0.18 |
